# A Single-Cell Level Comparison of Human Inner Ear Organoids and the Human Cochlea and Vestibular Organs

**DOI:** 10.1101/2022.09.28.509835

**Authors:** Wouter H. van der Valk, Edward S.A. van Beelen, Matthew R. Steinhart, Carl Nist-Lund, John C.M.J. de Groot, Peter Paul G. van Benthem, Karl R. Koehler, Heiko Locher

## Abstract

Genetic inner ear disorders are among the most common congenital abnormalities and lead to hearing loss and balance disorders. Ideally, tissue culture models of the inner ear should contain a functional unit combining otic sensory and nonsensory cell types to recapitulate the varied etiologies of inner ear disorders. Here, we evaluated cell type diversity of late-stage human pluripotent stem cell-derived inner ear organoids using single-cell transcriptomic analysis, electron microscopy and immunohistochemistry. We observed the induction of on-target inner ear-related periotic mesenchymal cells alongside off-target induction of skeletal myocytes, endothelial cells, and ependymal cells. By constructing a single-cell transcriptomic atlas of the human fetal and adult inner ear, we show that epithelium in the inner ear organoids contains cochlear and vestibular identities similar to the developing human inner ear. Moreover, the inner ear organoids contain immature type I and type II vestibular hair cells. Within these putative inner ear cell types, we confirmed the expression of genes and proteins linked to sensorineural hearing loss. This approach using human inner ear organoids would allow for disease modeling of specific genetic inner ear pathologies in the sensory and nonsensory domains of the inner ear.

## Introduction

The inner ear comprises the cochlea and vestibular organs, which mediate sound perception and balance. Functional impairment of the inner ear is common, with over 5% of the general population who suffers from hearing loss or balance disorders (Hülse et al., 2019; WHO, 2021). The etiology of these inner ear disorders varies greatly depending on age, genetic background, and environmental factors. Sensorineural hearing loss (SNHL) is one of the most common congenital disorders and impacts more than 1 in 1,000 newborns (Butcher et al., 2019; Sontag et al., 2020). Congenital inner ear disorders can be attributed to either syndromic or nonsyndromic genetic causes (Shearer et al., 1993; Van Beeck Calkoen et al., 2019). Based on the partial overlap of gene expression between the cochlea and the vestibular organs, it is not surprising that mutations in deafness genes may also lead to vestibular dysfunction, as seen in patients with DNFA9, DFNA11, DFNA15, and Usher syndrome (Mei et al., 2021).

Despite research efforts to understand the mechanisms of genetic SNHL and to identify gene therapies, a significant challenge is the heterogeneity of genetic SNHL (Taiber and Avraham, 2019). Current animal models are limited to modeling the effect of specific gene mutations. An alternative would be human inner ear cell cultures, which provide an avenue to evaluate the impact of SNHL genes on human inner ear development and function, as discussed in recent reviews (Czajkowski et al., 2019; Durán-Alonso, 2020; Roccio, 2021; Roccio and Edge, 2019; Roccio et al., 2020; Stojkovic et al., 2021; Tang et al., 2020; Zine et al., 2021). Human inner ear organoids (IEOs) for instance, can be derived from human pluripotent stem cells (hPSCs), either embryonic (hESCs) or induced (hiPSCs), using differentiation strategies that result in 2D or 3D cultures containing inner ear-like cells (Tang *et al*., 2020). Several SNHL gene mutations have been modeled in mouse and hPSC-derived otic culture systems (Van der Valk et al., 2021). However, most reports focus on hair cells, and it remains unclear to what extend PSC-based systems can be used to model congenital SNHL that impacts other cell populations of the inner ear (Van der Valk et al., 2021).

A promising system lies in the hPSC-derived IEOs (Koehler et al., 2017). The cellular diversity within these IEOs might be broader than previously reported, since hair-bearing skin organoids could be generated using a slightly modified IEO culture protocol (Lee et al., 2020). Moreover, recent and ongoing work on the temporal cellular diversity within the early IEO aggregates shows that this approach results in multilineage aggregates (Steinhart et al., 2021). Our goal in the current study was to determine the cellular diversity of late-stage hPSC-derived IEOs (differentiation day 75 and later), focusing on the inner ear cell types and comparison across cell lines. To this end, we provide a tool to determine early differentiation efficiency using multiple hPSC lines. We demonstrate on-target induction of inner ear epithelia, neurons, and periotic mesenchyme, but likewise identify off-target induction of epithelial, mesenchymal, endothelial, and neuro-ectodermal cell populations, including cranial skeletal myocytes, ependymal cells and vascular endothelial cells. For comparison, we generated an age-matched single-cell transcriptomic atlas of the fetal human inner ear. In addition, we include single-cell data from adult human vestibular organs to show the extent to which IEOs contain mature cell types of the human inner ear. Our comparative analysis reveals that organoids contain both sensory and nonsensory vestibular epithelial types, including type I and type II vestibular hair cells, and periotic mesenchyme. Surprisingly, we could identify a subset of cells resembling cochlear nonsensory epithelial cells. Furthermore, we assigned the expression of known SNHL or balance disorder-associated genes and proteins, which can be used as a resource for disease modeling study design and therapy testing in organoids.

## Results

### BMP signaling underlies variability in IEO induction across hPSC lines

We aimed to induce inner ear epithelium from multiple hPSCs using a streamlined method for IEO induction (Koehler et al., 2017; Steinhart et al., 2021) (Figure 1A). To generate IEOs, we enzymatically dissociated hPSCs into a single cell suspension and aggregated them into spheroids. After transferring the pluripotent spheroids two days later (differentiation day 0: D0), we exposed them to a low concentration of Matrigel, low concentration of basic fibroblast growth factor (bFGF), bone morphogenetic protein 4 (BMP-4) and SB-431542 to inhibit the transforming growth factor β (TGFβ) pathway and to induce a non-neural identity. bFGF and LDN-193189, a BMP inhibitor, were then added to induce an otic-epibranchial progenitor (OEPD) domain after three days (D3). To enhance otic induction, the Wnt pathway was activated by adding CHIR99021 at D8 and supplemented on D10. At D12, the aggregates were transferred to an ultra-low adhesion 24-well plate and allowed to self-organize into multi-organ aggregates. During this organization, otic placodes invaginate to form otic vesicles that mature to IEOs containing hair cells embedded in epithelial vesicles that receive neuronal projections.

**Figure 1.**
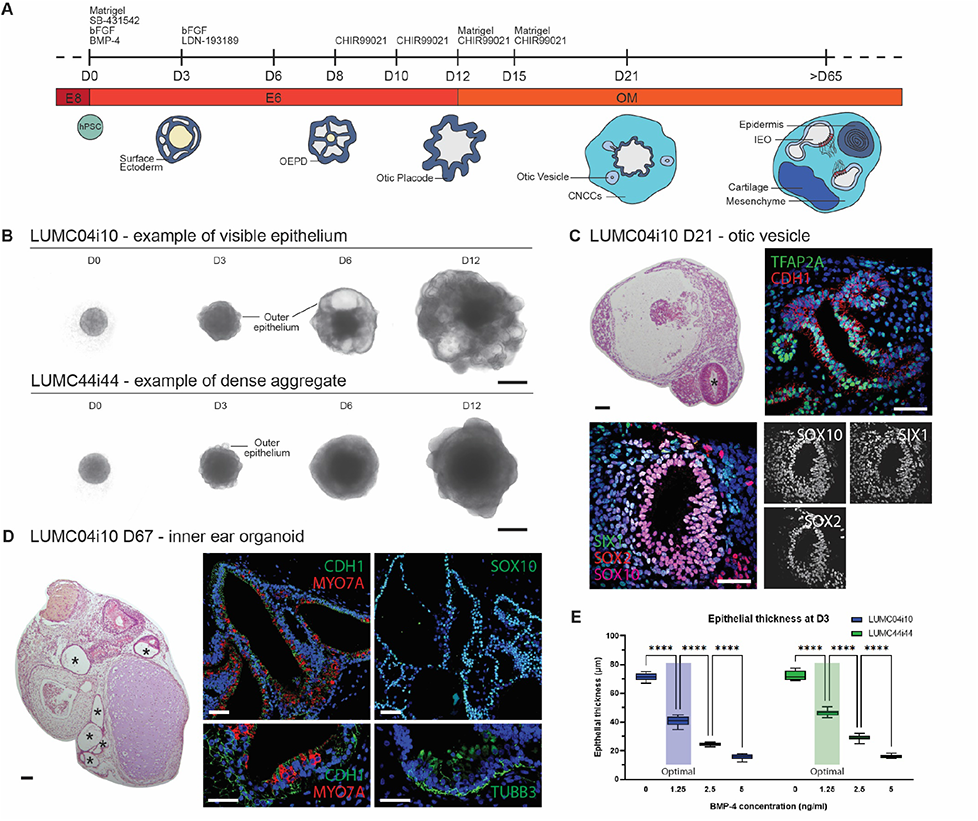
Otic induction efficiency is determined by BMP-4 concentration. **A.** Overview of the guided differentiation of inner ear organoids. **B.** Representative bright-field images of cell aggregates at D0, D3, D6 and D12 of the LUMC04i10 and LUCM44i44 hiPSCs with morphological differences between these lines from D6 onwards, containing either a visible epithelium (LUMC04i10) or a dense aggregate (LUMC44i44). Scale bars 500 μm. **C.** Representative HE and IHC images of D21 aggregates containing otic vesicles (asterisk in HE image) expressing TFAP2A, CDH1, SOX10, SIX1 and SOX2. Scale bar HE 100 μm, scale bars IHC 50 μm. **D.** HE and IHC images of D67 aggregates showing an IEO with epithelial vesicles, highlighted by CDH1 and SOX10, containing hair cells (MYO7A^+^) and neurons (TUBB3^+^). Asterisks mark IEO vesicles. Scale bar HE 100 μm, scale bars IHC 50 μm. **E.** Epithelial thickness measurements of D3 aggregates treated with different BMP-4 concentrations at D0 of the LUMC04i10 and LUMC44i44 cell lines. Efficient IEO induction was achieved with a BMP-4 concentration of 1.25 ng/ml (‘Optimal’). Statistical significance was analyzed with 2-way ANOVA with a Sidak correction for multiple comparisons. Data was considered statistically significant if P<0.05. n=10 per datapoint was graphed in a min-max box plot. A representative graph of 2 individual experiments is shown. CNCCs: cranial neural crest cells; D: differentiation day; IEO: inner ear organoid; OEPD: otic-epibranchial placode domain.

To evaluate the robustness of the differentiation protocol, we used eight human healthy-control hPSC lines: one hESC line (WA01) and seven hiPSC lines (WTC-SOX2, WTC-GCaMP, LUMC04i10, LUMC44i44, GON0515-03, GON0926-02, and SAH0047-02). In adapting the protocol to a broader cohort of cell lines, we noted that the initial BMP-4 treatment at D0 was a critical determinant of successful IEO production. In our experience, BMP-4 activity differs between vendors and lots, hampering consistent differentiation efficiency. We sought to determine which morphological features during early differentiation could be used to evaluate IEO induction efficiency objectively. To this end, we treated the cell lines with BMP-4 concentrations ranging from 0-20 ng/ml, and we followed morphologic features up to D12. During these first 12 days, we noticed a distinct variation in the morphology between cell lines (Figure 1B). LUMC04i10, WTC-SOX2, WTC-GCaMP, and SAH0047-02 are cell lines in which the developing outer epithelial layer can be clearly monitored over time until D12 (Figure 1B). This epithelial layer is composed of ectoderm (Koehler et al., 2017). LUMC44i44, WA01, GON0515-03, and GON0926-02 aggregates became denser after D3, and the outer epithelial layer could not be followed over time (Figure 1B). We, therefore, limited the measurement of morphological features to D3. Differentiation efficiency within these eight cell lines was determined by the presence of otic vesicles at D21 or the presence of more mature IEOs at D65 and later, using hematoxylin and eosin (HE) staining or immunohistochemistry (IHC) (Figure 1C-D). The otic identity of these inner ear vesicles could be confirmed by the expression of CDH1, TFAP2A, SOX10, SOX2, and SIX1 (Figure 1C). The IEOs of D65 and later were CDH1^+^ SOX10^+^, contained MYO7A^+^ hair cells with TUBB3^+^ neurons that protrude towards hair cells (Figure 1D). The highest differentiation efficiency was reached at a BMP-4 concentration of 1.25 ng/ml for this set of experiments using BMP-4 from the same stock, with either otic vesicles or epithelial IEOs forming in all cell lines, independent of their variation in morphology beyond D3. We measured epithelial thickness, circularity, and size of the aggregates *in vitro* at D3 (Figure S1A). Epithelial thickness is hypothetically useful, as the ectoderm decreases in thickness from medial to lateral (Figure S1B) (Mayordomo et al., 1998) based on BMP activity levels (Groves and Fekete, 2012; Wilson and Hemmati-Brivanlou, 1995). Circularity and size have been described to alter during IEO differentiation and potentially also differ between BMP-4 concentrations (Koehler et al., 2017). From these morphological features, only epithelial thickness significantly differed between BMP-4 concentrations over all hPSC lines. Thickness decreased at the highest BMP-4 concentrations (Figure 1D, Figure S1B-C). Efficient IEO differentiation was achieved with aggregates containing an outer epithelium layer with on average a thickness of 35.9 µm (S.D. 6.2 µm, n=10 aggregates per cell line; 8 cell lines). Size and circularity of the aggregate were also influenced by the BMP-4 concentration, however less consistent over all cell lines (Figure S1C). Depending on the BMP-4 lot number or vendor however, BMP-4 concentrations up to 20 ng/ml had to be used to reach the same epithelial thickness and subsequent efficient induction of otic vesicles and IEOs (data not shown). In conclusion, optimal BMP-4 concentration at D0 can be inferred from morphological changes as early as D3 and not necessarily by the absolute concentration of BMP-4. A range of 1.25-20 mg/ml is acceptable depending on the vendor, handling, and storage (see the Material and Methods section for guidance on vendor selection and handling). Using the optimal BMP-4 concentration, we efficiently induced otic vesicles in all hPSC lines tested.

### Single-cell and single-nucleus RNA sequencing unravel the cellular diversity of IEOs

To comprehensively characterize the cell types that arise in the efficiently differentiated aggregates, we performed single-cell RNA sequencing (scRNAseq) as well as single-nucleus RNA sequencing (snRNAseq) on D75 to D110 aggregates derived from four hPSC lines (WA01, WTC-SOX2, WTC-GCaMP, and LUMC04i10). The WTC-SOX2 and WTC-GCaMP lines are commonly used for our experiments and therefore included, although their reporter functions are beyond the scope of this manuscript. We decided to include snRNAseq in addition to scRNAseq as certain cell types, such as epithelial cells, might be better presented by the single nucleus approach (Korrapati et al., 2019; Slyper et al., 2020). More importantly, both methods allow for similar discrimination of cell types when intronic sequences are included in snRNAseq analyses (Bakken et al., 2018; Ding et al., 2020). In addition to dissociation of wholes aggregate into single cells, we used another approach by which aggregates were first cut in 200 µm thick slices using a vibratome. This approach allowed us to select sections that contain IEO-like vesicles by bright-field evaluation before dissociation into single cells.

The cellular heterogeneity of the aggregates is both captured in the scRNAseq (Figure S2A-B) and snRNAseq data (Figure S3A-B). The subtypes were annotated using expression of cell type-specific and marker genes (Figure S2C-D; Figure S3C-D). With the scRNAseq approach, we identified the same cell types for both the WTC-SOX2 and WTC-GCaMP lines, at D75 and D100, irrespective of whether the aggregates were vibratomed or not. The vibratome approach, however, did not result in a relative larger number of otic cell types (Figure S2B). With the snRNAseq approach, the same cell types are captured for both the WA01 and LUMC04i10 lines, at D75 and D110. In these experiments, both the WA01 and LUMC04i10 cell lines were combined after dissociation into single nuclei. We used cell line-unique single nucleotide polymorphisms (SNPs) to demultiplex the data, by which we could assign a cell line identity to 97.3% of the nuclei. We identified some variation between the scRNAseq and snRNAseq approach: using scRNAseq of the WTC-SOX2 and WTC-GCaMP hiPSC lines, clusters of cycling cells and cells with a high ribosomal content were identified that could not identified using the snRNAseq approach in the WA01 and LUMC04i10 hiPSC lines. Similarly, adipocytes could not be identified using scRNAseq of the WTC-SOX2 and WTC-GCaMP hiPSC lines, but were seen in the snRNAseq approach.

As we were able to identify similar cell types with both approaches, we integrated the data to a combined dataset of sequenced cells and nuclei for further analyses (Figure 2A-C; Figure S4). Of these cell types in the aggregate, 8.3% (3.7% to 11.1% between cell lines) assigned to an otic identity that includes otic epithelium, hair cells or periotic mesenchyme (POM) (Figure 2C). We wondered if the other populations contained cells reminiscent of the inner ear spatial environment, thus we set out to analyze these populations. Most of these cell types were of mesenchymal origin (68.0%, Figure 2B). This group contained a cluster of myocytes (*TNNC1*, *TPM3*, *SRL*, *CKM*, *LDB3*) (Lindskog et al., 2015) (Figure 2D). The presence of these myocytes corresponds with our observations of irregular and sporadic contractions of later-stage aggregates *in vitro* (data not shown). This mesenchymal population expressed skeletal myocyte-specific genes (*TNNT1*, *MYBPC1*, *MYOT*, *RYR1*, *JPH1*) (Figure S5A) (Lindskog et al., 2015), with a marked lack of expression of marker genes for either smooth muscle cells (*ACTA2*, *MYOCD*, *MYH11*) (Dobnikar et al., 2018) or cardiomyocytes (*MYH6*, *MYL7*, *TNNI3*, *MYBPC3*, *CACNA1C*) (Lindskog et al., 2015). In line with the fetal skeletal myocyte gene expression of *MYL4*, *ACTC1* and *TNNT2* (Figure S5A) (Anderson et al., 1991; Vandekerckhove et al., 1986; Whalen et al., 1978), we could identify different transcriptional stages of skeletal myogenic differentiation within this population (Figure 5SB): myocyte precursors (*PAX7*), myoblasts/myocytes (*MYOD1*,*MYOG*), and myotubes (*MYL2*, *MYH2*, *MYH7*, *ITBG1*) (Choi et al., 2020). Interestingly, when identifying upstream genetic regulators that control cranial myogenesis, we observed *TBX1* and *PITX2* gene expression, with a relative absence of *PAX3* (Figure S5C), limiting the possible identity of these skeletal myocytes to that of cranial skeletal myocytes, as reviewed in (Grimaldi and Tajbakhsh, 2021). Moreover, moderate expression of *NKX2-5* could be seen in this cluster (data not shown). Although *NKX2-5* expression is well established in cardiomyocyte development, expression on the protein and mRNA level has been described in developing cranial skeletal myocytes (Kasahara et al., 1998). Using IHC, we confirmed the presence of ACTC1^+^ TNNT1^+^ MYOG^+^ skeletal myocytes in our aggregates (Figure 2E). The mesenchymal population consists also of a putative chondrocyte population (*MATN4*, *COL9A1*, *SOX6*, *SOX9*, *COL2A1*) which contains an early mature signature (*CNMD*, *EPYC, FRZB*) (Figure S5D) with more mature genes such as *ISG15*, *IFI6* and *MX1* expressed only in a fraction of the cells (Wu et al., 2021) (Figure S5E). Additionally, we identified fibroblasts (*COL1A1*, *PDGFRA*, *TWIST2*, *APCDD1*, *DPT*) and pericytes (*CSPG4*, *PDGFRB*, *RGS5*, *KCNJ8*, *ABCC9*) forming distinct clusters within the mesenchymal cell types (Figure S5F-G).

**Figure 2.**
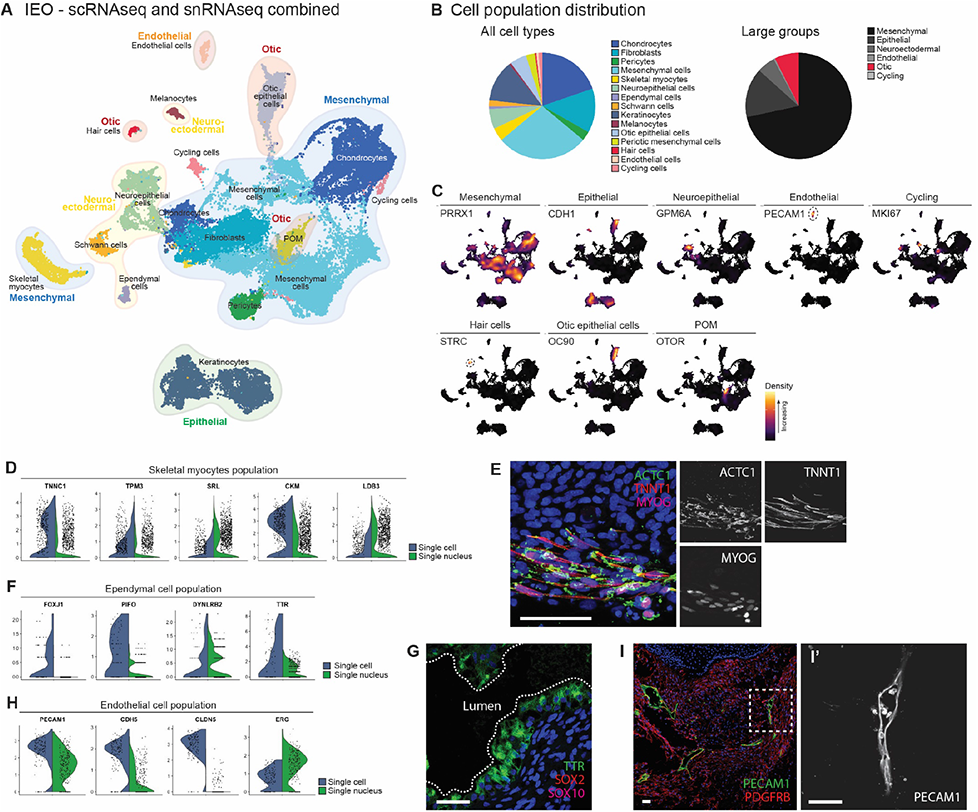
Single-cell transcriptomic analysis of D75-D110 aggregates reveals cellular diversity of IEOs. **A.** Overview UMAP plot of the combined scRNAseq and snRNAseq data of D75 to D110 IEOs with cell type annotations. **B.** Relative contribution of cell types and large groups. **C.** Marker genes involved in cell type annotation assignment. **D.** Marker analysis highlights expression of skeletal myocyte marker genes within the skeletal myocyte cluster. scRNAseq and snRNAseq are plotted separately. **E.** ACTC1^+^ TNNT1^+^ MYOG^+^ skeletal myocytes in a D75 aggregate. Scale bars 50 μm. **F.** Ependymal marker gene expression in the ependymal cell cluster. scRNAseq and snRNAseq are plotted separately. **G.** TTR^+^ SOX2^-^ SOX10^-^ ependymal cells in a D75 aggregate. Scale bar 50 μm. **F.** Endothelial marker genes within the endothelial cluster. scRNAseq and snRNAseq are plotted separately. **I.** PECAM1^+^ endothelial cells forming luminal-like structures within a D75 aggregate, at high magnification (I’). Scale bars 50 μm. POM: periotic mesenchyme.

The second largest group within the aggregate are the epithelial cells consisting of keratinocytes and otic epithelial cells (12.6%). The keratinocytes within the aggregate showed a similar cell type-diversity to the skin organoids (Lee et al., 2020). Within this population expressing *CDH1*, *KRT5* and *KRT19*, we could identify basal (*CXCL14*), intermediate (*KRT1*) and peridermal (*KRT4*) keratinocytes (Figure S5H).

Neuroectodermal cell types are involved in certain types of SNHL, such as Waardenburg syndrome affecting the melanocyte lineage, but also in neurofibromatosis type II, which is characterized by neoplastic Schwann cells (Truong et al., 2018). To get a better understanding of these populations, we set out to determine how mature these cell types are in our IEOs. The neuroectodermal group (9.4%) consisted of neuroepithelial cells, ependymal cells as well as Schwann cells and melanocytes. The neuroepithelial population could be divided into a group of neuroepithelial progenitor cells (*SOX2*, *SNHG11*) and more mature central (*GPM6A*, *MATK*, *RASFRF1*) and peripheral (*PRPH*, *PIRT*) neurons (Figure S5I) (Zeisel et al., 2018). These latter expressed genes for ion channels known to be enriched in the peripheral nervous system and more specifically in sensory neurons (*P2RX3*, *SCN7A*, *SCN10A*) (Wangzhou et al., 2021). When analyzing semi-thin sections, we identified occasional groups of neurons in the stroma surrounding the IEOs. They contain a distinct cell body with a large, round nucleus and a single dendritic process, suggestive for neurons (Figure S5J). The dendritic processes were surrounded by cells with elongated, flat nuclei with morphologies similar to Schwann cells. Using transmission electron microscopy (TEM), we observed evidence of Schwann cells associated with these neurons near the otic vesicles within the IEOs (Figure S5K). The presence of these Schwann cells was confirmed by the transcriptomics data, with a putative Schwann cell population expressing marker genes (*SOX10*, *ERBB3*, *S100B*, *PLP1*), as well as genes for immature (*NGFR*, *NCAM1*, *L1CAM*, *CDH2*) and (pro)-myelinating Schwann cells (*POU3F1*, *CDKN1C*, *MPZ*, *PTN*, *CRYAB*) (Figure S5L). We also discovered a population of ependymal cells that expressed *FOXJ1*, *PIFO*, *DYNLRB2* and *TTR* (Figure 2F) (MacDonald et al., 2021). We confirmed the presence of TTR^+^ SOX2^-^ SOX10^-^ ependymal cells forming vesicle-like structures within the aggregate (Figure 2G), in contrast to the TTR^-^ SOX2^+^ SOX10^+^ otic vesicles. The last neuroectodermal population was composed of melanocytes, which expressed genes of various stages of melanocyte differentiation, including marker genes for melanoblasts (*SOX10*, *EDNRB*, *DCT*, *MITF*), melanocytes (*KIT*, *PAX3*, *EDNRB*) and that of differentiated fully pigmented melanocytes (*TYRP1*, *TYR*, *GPR143*) (Figure S5M) (Belote et al., 2021; White and Zon, 2008). Although melanocytes are generally known for their role in skin pigmentation (Lee et al., 2020), a subset of melanocytes incorporates into the ionic regulatory epithelium in the inner ear (Locher et al., 2015; Van Beelen et al., 2020); hence, we sought to identify this population in IEOs. Interestingly, melanocytes (MLANA^+^) were found within or very closely related to the SOX10^+^ IEOs (Figure S5N), similar to the developing human fetal vestibular organs (around fetal week 10) and cochlea (around fetal week 11) (Locher et al., 2015; Van Beelen *et al*., 2020).

We discovered a population of endothelial cells in the IEOs using the transcriptomic data that could be reminiscent to the blood-labyrinth barrier of the inner ear. This population (0.7%) expressed known endothelial markers *PECAM1*, *CDH5*, *CLDN5*, and *ERG* (Figure 2H) (Schupp et al., 2021), with concomitant expression of vascular marker genes (*FLT1*, *VWF*, *NOTCH4*, *NRP1*) (Berendam et al., 2019; Schupp et al., 2021) and relative absence of lymphatic marker genes (*PROX1*, *PDPN*) (Figure S5O) (Kalucka et al., 2020). *LYVE1* and *FLT4*, both expressed in this population, are known lymphatic-specific genes, although these genes have been observed during fetal vascular endothelial development (Gordon et al., 2008; Kaipainen et al., 1995). We could confirm the presence of PECAM1^+^ endothelial cells within the aggregate forming structures that contain a lumen throughout the aggregate (Figure 2I).

In summary, using different hPSC lines, we could robustly induce multilineage cell types within the IEOs. The cell types we identified are known to be expressed in the environment of the inner ear, although we could not grant all of them with an “inner ear identity”. In addition to the putative cranial identity of the skeletal myocytes, we identified gene expression profiles that match fetal or embryonic development, for instance for the skeletal myocytes, chondrocytes, and endothelial cells. All cell types are represented using any cell line, at any timepoint and using either scRNAseq or snRNAseq (Figure S4B), although, there were differences in relative cell type contribution between experiments. WTC-GCaMP and WTC-SOX2 aggregates (which share a parental genome) showed a similar cell type distribution based on scRNAseq data. However, this distribution differed from the LUMC04i10 and WA01 cell line that are used for snRNAseq. This could be due to variability in dissociation method (either single cell or nucleus), or by cell line variation. The latter is supported by the difference we observe between the WA01 and LUMC04i10 cell line, notably the large neuroepithelial cell type cluster found in the WA01 cell line and a relatively smaller mesenchymal cluster. In this manuscript, we did not perform thorough batch-to-batch variation tests to rule out the role of cell lines or the transcriptomics method used.

### Generation of an atlas of fetal and adult human inner ear tissues

To compare cell type diversity within IEOs to those found in the native human inner ear, we carried out snRNAseq on fresh inner ear tissue collected at different stages of human inner ear development. We collected one fetal inner ear of fetal age week 7.5 (W7.5), one fetal inner ear of W9.2, and an ampulla and utricle from a 47-year-old donor (Figure 3A). At W9.2, we would expect that most of the mature cochlear cell types are still in development and the epithelial populations could be separated out in different domains (Figure 3B). The cochlear duct floor divides in a medial, prosensory and lateral domain, giving rise to the future Kölliker’s organ, organ of Corti and outer sulcus, respectively. In addition, the roof of the cochlear duct will give rise to Reissner’s membrane and the stria vascularis, reviewed in (Driver and Kelley, 2020). Vestibular development, which precedes that of the cochlea, has a more mature phenotype distinct epithelial cell types at W9.2 (Johnson Chacko et al., 2016; Van Beelen *et al*., 2020).

**Figure 3.**
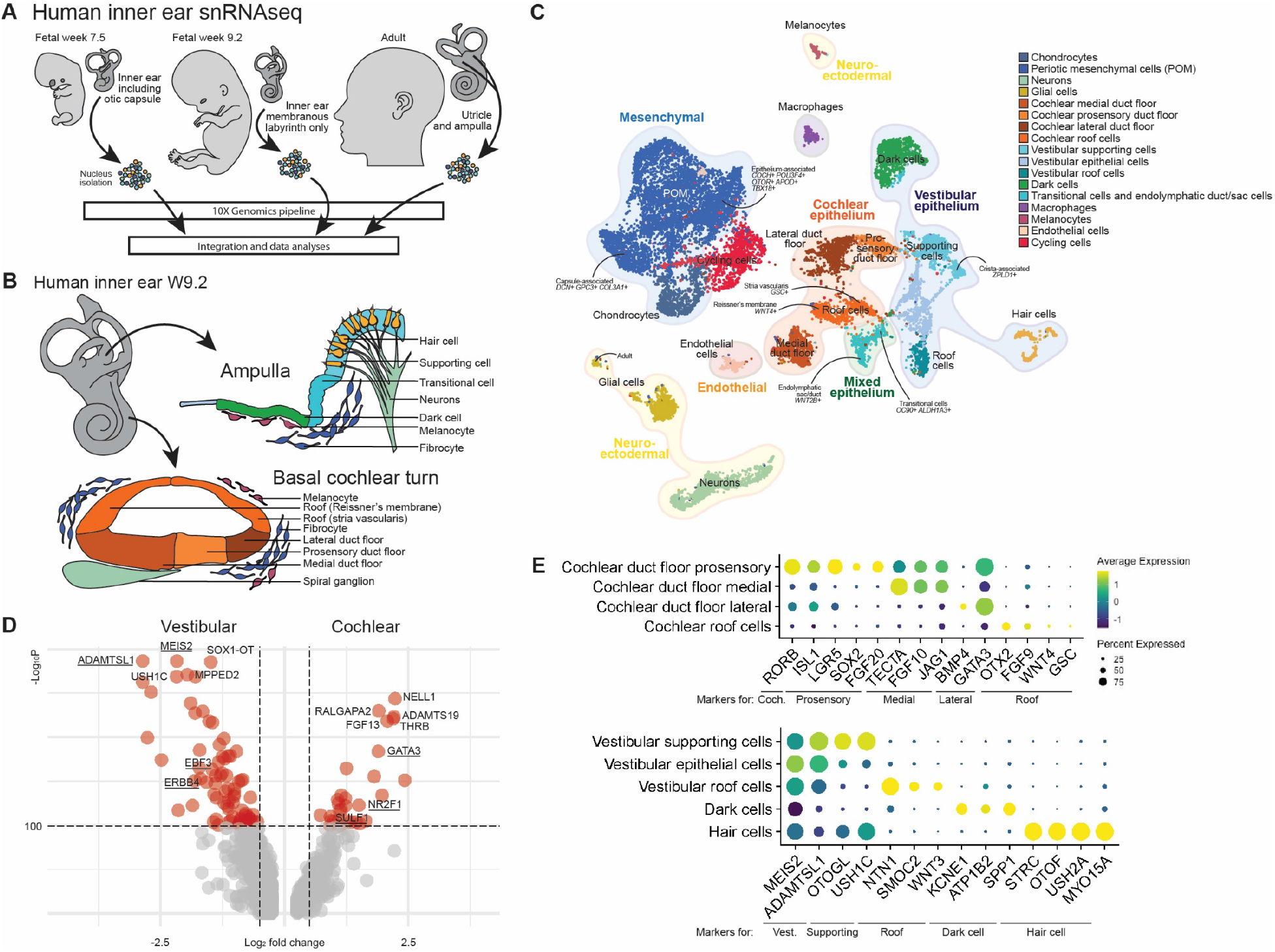
Generation of a human fetal and adult inner ear single-cell atlas. **A.** Experimental overview of the collection of fresh human inner ear tissue from W7.5, W9.2 and an adult donor. **B.** Schematic of the human inner ear at W9.2 of development, depicting the cellular diversity of the ampulla as example of the vestibular organs and the basal turn of the cochlea. **C.** Overview UMAP plot of the combined human W7.5, W9.2 and adult inner ear dataset with cell type annotations. **D.** Volcano plot showing differentially expressed gene between the vestibular supporting cells and cochlear duct floor (medial, prosensory and lateral). Underlined genes have recently been described to be differentially expressed between vestibular and cochlear cell types. **E.** Dotplot showing expression of known marker genes within cochlear and vestibular domains and specific cell types. Coch.: Cochlear; Vest.: Vestibular.

We used recently published scRNAseq and snRNAseq datasets of the mouse inner ear (Jan et al., 2021; Korrapati et al., 2019; Wilkerson et al., 2021; Yamamoto et al., 2021; Yu et al., 2019) to annotate the populations of the integrated fetal W7.5, W9.2 and adult inner ear datasets (Figure 3C). We identified a mesenchymal population, various epithelial populations of both cochlear and vestibular identity, hair cells, neurons, glial cells, cycling cells, endothelial cells, macrophages, and melanocytes (Figure 3C; Figure S6). To confirm the vestibular versus the cochlear identity of the epithelial cell population, we analyzed the differential gene expression between the putative cochlear duct floor and vestibular supporting cells (Figure 3D). *ADAMTSL1*, *MEIS2*, *EBF3* and *ERBB4* were genes highly expressed within vestibular populations, in contrast to *GATA3*, *NR2F1* and *SULF1* which are present in cochlear populations, as has been shown on a single-cell level for the developing mouse inner ear (Wilkerson et al., 2021; Yamamoto et al., 2021). The cochlear cell types could be separated (Figure 3E) into a medial (*TECTA*, *FGF10*, *JAG1*), prosensory (*ISL1*, *LGR5*, *SOX2*, *FGF20*) and lateral domain of the duct floor (*BMP4*, *GATA3*) as well as the cochlear roof (*OTX2*, *FGF9*, *WNT4*, *GSC*), based on data from the developing mouse cochlea (Chai et al., 2011; Hayashi et al., 2008; Luo et al., 2013; Morsli et al., 1999; Munnamalai and Fekete, 2020; Ohyama et al., 2010; Yamamoto et al., 2021). The vestibular population is composed of putative supporting cells (*OTOGL*, *USH1C*) with a crista-associated *ZPLD1*^+^ supporting cell population, roof cells (*NTN1*, *SMOC2*, *WNT3*) and dark cells (*KCNE1*, *ATP1B2*, *SPP1*) expressing marker genes that have recently been described for the developing mouse vestibular organs (Jan et al., 2021; Wilkerson et al., 2021). The remaining epithelial cluster was composed of a mixed population of endolymphatic duct/sac cells (*WNT2B*) and vestibular transitional cells (*OC90*, *ALDH1A3*) (Honda et al., 2017; Wilkerson et al., 2021).

Within the mesenchymal population, chondrocytes and a large cluster of periotic mesenchymal cells (POM) were identified. The chondrocytes were mostly identified in the W7.5 inner ear, as during preparation of this sample the otic capsule was not removed from the membranous labyrinth. As recently described for the developing mouse vestibular organs, different cell populations could be identified in the mesenchymal population, with a transcriptomic difference between epithelium-associated mesenchyme and capsule-associated mesenchyme (Wilkerson et al., 2021). We observed a similar diversity in this dataset, with capsule-associated POM (*DCN*, *GPC3*, *COL3A1*) clustering closely to the chondrocytes and distinctly separate from the epithelium-associated POM (*COCH*, *POU3F4*, *OTOR*, *APOD*, *TBX18*). To the best of our knowledge, these data constitute the only single-cell level analysis of early-stage inner ear development combined with human adult vestibular organs and should provide a valuable resource to the community beyond their use for organoid comparative analysis.

### POM within the IEOs is associated with the sensory epithelium

To assess how the IEOs resemble the human inner ear, we used Symphony, a tool for single-cell reference mapping (Kang et al., 2021). Using this R package, IEO-derived POM, otic epithelium and hair cells were mapped as a query to the reference atlas of the human inner ear. We performed these analyses separate for the mesenchymal, epithelial and hair cell populations.

We mapped the putative POM cluster present in the IEO dataset to the human inner ear atlas reference (Figure 4A-B). A large overlap was present with the reference POM cluster, more specifically with the epithelium-associated POM. Expression of *CRYM*, *OTOR*, *COCH* and *POU3F4* within the IEO-derived POM confirmed the epithelium-associated identity (Figure 4C). The other region overlapping was composed of POM that was found in the human fetal inner ears (Figure S6B). We did not observe any overlap with the capsule-associated POM from the reference human inner ear atlas. To confirm the presence of the epithelium-associated POM, we performed additional IHC analyses of D75 IEOs. Surrounding inner ear vesicles containing MYO7A^+^ hair cells, we noticed the presence of OTOR^+^ POU3F4^+^ POM (Figure 4D). These OTOR-expressing cells resembled the OTOR^+^ mesenchymal cells *in vivo* that directly underly the sensory epithelium of the fetal human utricle (Figure 4E).

**Figure 4.**
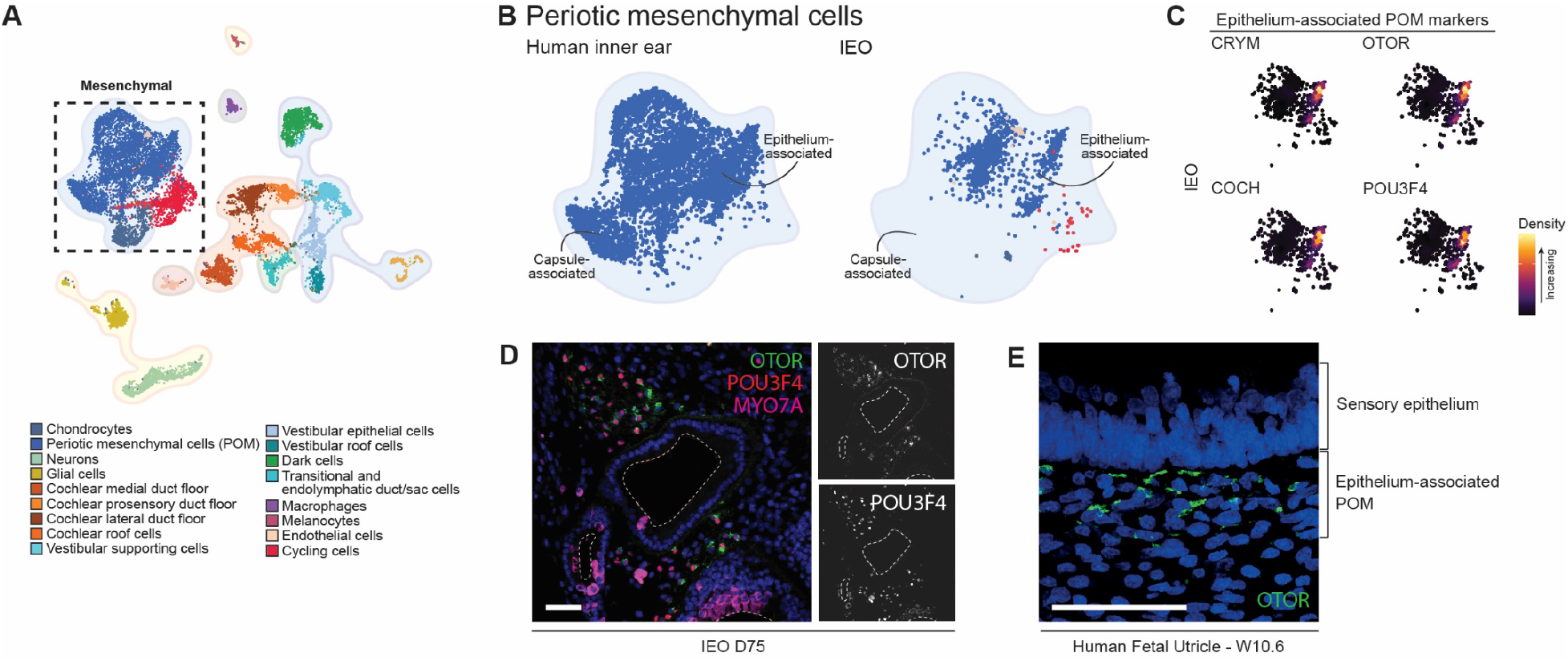
IEO-derived POM resembles human fetal epithelium-associated POM. **A.** Overview UMAP plot of the annotated human inner ear atlas with the mesenchymal cell types highlighted. **B.** UMAP plot of the reference POM (left) and mapped IEO-derived POM population (right) as analyzed by Symphony. **C.** Density plots of the IEO-derived POM population showing expression of epithelium-associated markers. **D.** OTOR^+^ POU3F4^+^ mesenchymal cells surrounding IEO-vesicles containing MYO7A^+^ hair cells at D75. Scale bar 50 μm. **E.** OTOR^+^ mesenchymal cells directly underlying the sensory epithelium of the fetal utricle (W10.6). Scale bar 20 μm.

### IEOs contain immature type I and type II vestibular hair cells

IEO-derived hair cells were mapped to hair cells of the human inner ear atlas reference using Symphony (Figure 5A). We do not expect that any cochlear hair cells were captured in the human inner ear atlas, as these arise only by W10 of fetal development (Locher et al., 2013; Pujol and Lavigne-Rebillard, 1985). Within the reference hair cells, immature developing hair cells clustered separately from the adult hair cells (Figure 5B; Figure S6B). Immature hair cells expressed *SOX2*, *ATOH1* and *ANXA4* (Figure 5C). Two distinct adult hair cell populations were present, showing specific expression for markers of type I and type II vestibular hair cells, *OCM* (Simmons et al., 2010) and *ANXA4* (McInturff et al., 2018), respectively (Figure 5C). The IEO-derived hair cells overlapped with the developing immature hair cells and expressed *SOX2* and *ATOH1* (Figure 5B-C).

**Figure 5.**
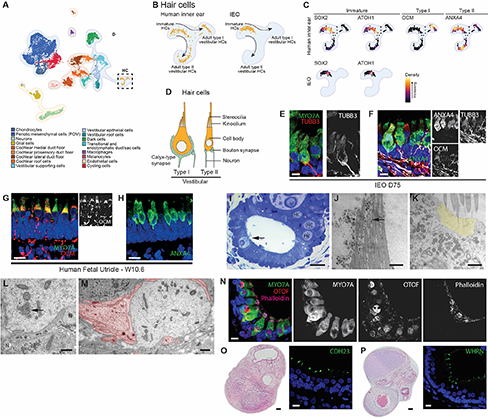
Immature type I and type II vestibular hair cells are present in the IEOs. **A.** Overview UMAP plot of the annotated human inner ear atlas with the hair cells (HC) highlighted. **B.** UMAP plot of the reference hair cell population (left) and mapped IEO-derived hair cell population (right) as analyzed by Symphony. **C.** Density plots of the reference hair cell population (left) for markers of immature, type I and type II vestibular hair cells, as well as density plots for the mapped IEO-derived hair cell population showing expression of immature markers. **D.** Schematic of type I and II vestibular hair cells depicting their morphological differences. **E.** Calyx-type synapse (TUBB3A^+^) surrounding a hair cell (MYO7A^+^). Scale bar 10 μm. **F.** Expression of ANXA4 and OCM in hair cells of a D75 IEO, showing a subset of hair cells expressing none, either or both. Scale bar 10 μm. **G.** OCM expression in hair cells in the human fetal utricle (W10.6). Scale bar 10 μm. **H.** ANXA4 expression in the human fetal utricle (W10.6). Scale bar 10 μm. **I.** Staining (methylene blue-azure II) of a D74 IEO-vesicle containing hair cells (arrow and “HC”). Scale bar 20 μm. **J.** Hair bundle comprising the stereocilia and (part of) the kinocilium. Arrow points toward cross-section of the kinocilium. Scale bar 1 μm. **K.** Apical region of a hair cell showing its cuticular plate (yellow) with the stereociliary rootlets. Scale bar 1 μm. **L.** Basal region from a hair cell showing multiple synaptic ribbons (arrow). Scale bar 500 nm. **M.** Transection of a type I vestibular hair cell which is almost completely surrounded by the calyx-type synapse (in red). Scale bar 500 nm. **N.** OTOF expression in MYO7A^+^ hair cells. Scale bar 10 μm. **O.** HE overview with highlighted area showing CDH23 in stereocilia by IHC. Scale bar HE 100 μm, scale bars IHC 20 μm. **P.** HE overview with highlighted area showing WHRN in stereocilia by IHC. Scale bar HE 100 μm, scale bars IHC 20 μm. HC: hair cell.

Although type I and type II vestibular hair cells did not form distinct clusters in the reference immature hair cell cluster, a distinction can be made between developing type I and type II vestibular hair cells at W9 (Johnson Chacko et al., 2016). Type I and type II vestibular hair cells can be distinguished by their shape and the type of synapse (Figure 5D). Calyx-type synapses (TUBB3^+^) which are typical for type I vestibular hair cells, could be identified in D75 IEOs (Figure 5E). Additionally, within IEOs we could identify ANXA4^+^ hair cells, a known marker for developing hair cells and type II vestibular hair cells, with some of these hair cells also expressing OCM, a marker for type I vestibular hair cells (Figure 5F). This expression is similar to the human fetal utricle (Figure 5G-H). Additionally, we performed light-microscopical and ultrastructural analyses of the hair cells present in IEOs. The IEOs consisted of a central, open lumen lined by a single layer of columnar cells with small, densely stained nuclei located in their basal part (Figure 5I). Interspersed between these epithelial cells are less azurophilic, flask-shaped cells with a large, round nucleus and from which a large hair bundle protrudes into the lumen. The morphology of these cells closely resembles that of vestibular hair cells. Using TEM, the flask-like cells contain a large hair bundle, in which the individual stereocilia are of varying length (Figure 5J). These stereocilia are inserted into the cuticular plate in the apical part of the hair cell as is evident by the presence of stereociliary rootlets (Figure 5K). At the opposite pole of these hair cells, frequently seen features were synaptic ribbons which are oriented opposite the bouton-type synapses associated with the basal membranes of the hair cells (Figure 5L). As these synapses did not contain any vesicles it may be surmised that these are afferent neurons. The observation that these cells were associated with bouton-type synapses seemed to indicate that they were type II vestibular hair cells (Koehler et al., 2017). Other hair cells, however, resembled the calyx-type synapse seen with type I vestibular hair cells (Figure 5M).

Further analyses revealed that these immature vestibular hair cells express proteins of genes linked to SNHL. In addition to MYO7A (DFNA11, DFNB2, and Usher syndrome type I), OTOF (DFNB9), CDH23 (DFNB12 and Usher syndrome type I), and WHRN (DFNB31) are expressed within the hair cell or its stereocilia.

### Cochlear and endolymphatic duct/sac epithelium is induced in IEOs

Based on previous work, the IEO are assumed to have a vestibular sensory epithelium containing supporting cells (Koehler et al., 2017). In line with this hypothesis, most IEO-epithelial populations were mapped to the vestibular supporting cells and epithelial cells, with a total lack of vestibular dark cells and roof cells (Figure 6A-B). However, IEO-derived epithelial populations also overlapped with the endolymphatic duct/sac cells, expressing *WNT2B* (Figure 6C). Furthermore, a large overlap is seen with the cochlear medial duct floor, to a lesser degree for the cochlear lateral duct floor, where it is almost absent for the prosensory domain and cochlear roof cells. Marker gene expression of *TECTA* confirmed the medial duct floor-like population within the IEOs (Figure 6C). *TECTA* expression in the vestibular supporting cells may be explained by its temporal expression in the vestibular organs (Rau et al., 1999). The presence of these populations was consistent over cell lines and timepoints, although in varying degrees (Figure 6D).

**Figure 6.**
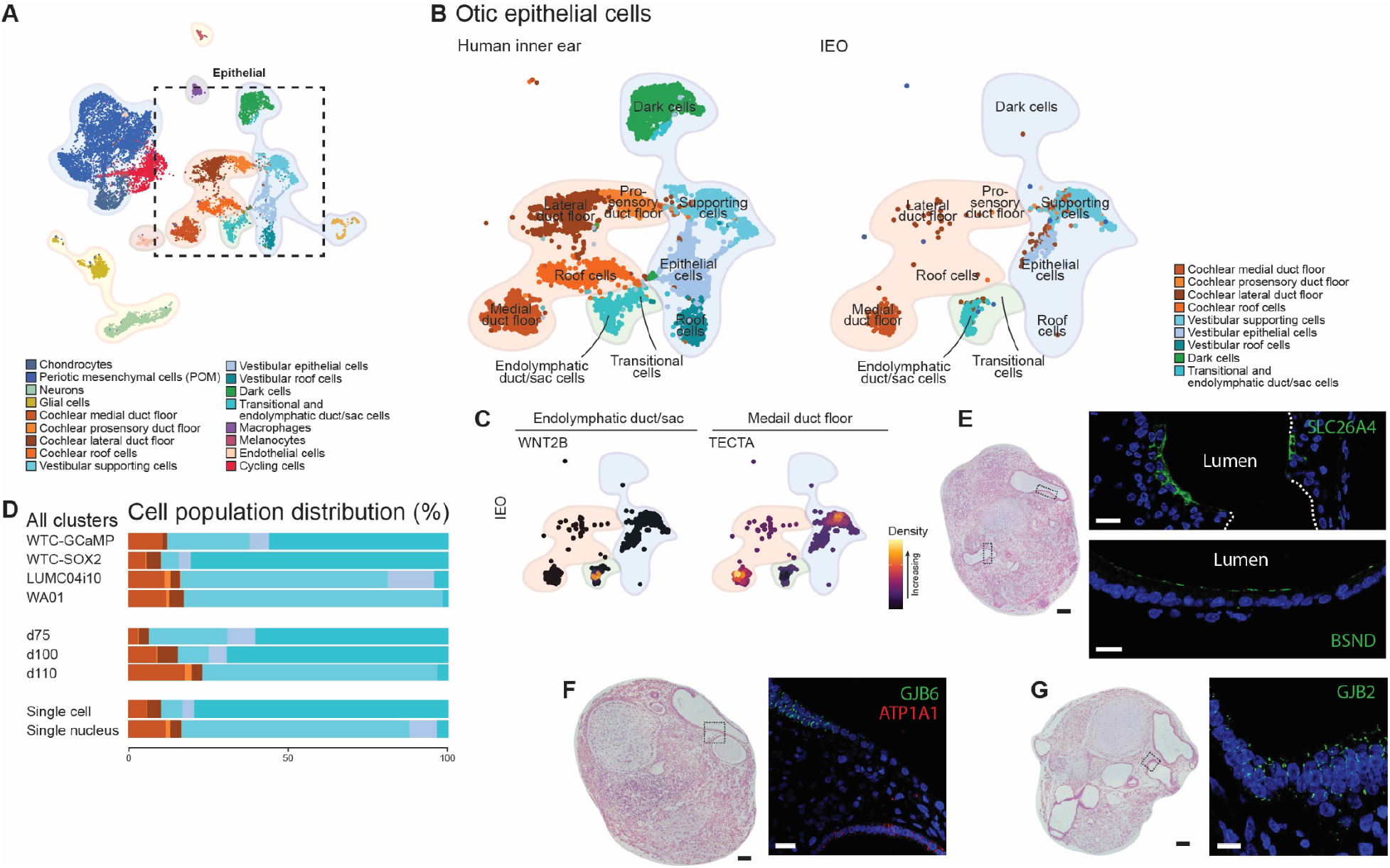
Endolymphatic duct/sac and cochlear epithelia in addition to vestibular supporting cells develop in IEOs. **A.** Overview UMAP plot of the annotated human inner ear atlas with the epithelial cell types highlighted. **B.** UMAP plot of the reference epithelial cell populations (left) and mapped IEO-derived otic epithelium population as analyzed by Symphony. **C.** Density plots of the IEO-derived epithelial populations showing expression of endolymphatic duct/sac marker *WNT2B* and medial duct floor marker *TECTA*. **D.** Relative contribution of cell types per line, timepoint and technique used. **E.** HE overview with in the highlighted area expression of SLC26A4 and BSND (D75). Scale bar HE 100 μm, scale bars IHC 20 μm. **F.** HE overview and localized expression of GJB6 and ATP1A1 within the highlighted area (D75 IEO). Scale bar HE 100 μm, scale bar IHC 20 μm. **G.** HE overview and expression of GJB2 within the highlighted epithelium (D75 IEO). Scale bar HE 100 μm, scale bar IHC 20 μm.

Using IHC, we were able to show expression of SLC26A4 and BSND, known to be expressed in specific epithelia of the inner ear including the endolymphatic sac (Honda et al., 2017). Mutations in *SLC26A4* may cause SHNL in DFNB4 and Pendred syndrome, and mutations in *BSND* leading to DFNB73. Within the IEO epithelia we were able to show additional expression of ATP1A1, GJB6, and GJB2 (Figure 6F-G). Mutations in *GJB2* are the most common cause of nonsyndromic SNHL causing DFNA3A and DFNB1A. Within epithelia these connexin-coding genes were expressed together with GJB6 that causes DFNA3B. These proteins are *in vivo* expressed in a spatiotemporal manner in the inner ear epithelia, including the human medial cochlear duct floor (Locher et al., 2015) and sensory vestibular epithelia (Forge et al., 2003). Moreover, ATP1A1, implicated in age-related hearing loss (Ding et al., 2018), has a spatiotemporal expression during development of the inner ear (Locher et al., 2013; Van Beelen et al., 2020), and shows epithelial expression within IEOs (Figure 6F).

### IEO express SNHL genes and proteins throughout the aggregate

A promising use of IEOs is disease modeling, including modeling of syndromic and nonsyndromic SNHL, which has been linked to over 150 genes. We leveraged our datasets to assess whether these genes are expressed within the proper cell subtypes of the IEO model. Most SNHL genes can be linked to the function of the protein in specific inner ear cell types (Delmaghani and El-Amraoui, 2020; Nishio et al., 2015). We compiled a comprehensive summary table of the gene/protein expression for 158 genes from the clinical literature (Table S1). By assessing the sparseness of gene expression in heatmaps and – for some genes – protein expression, we developed a scoring system to aid the reader in assessing whether a particular gene of interest could be studied using the IEO system (Table S1, Figure S7A-E). In total, 67 of 158 genes showed expression in IEOs in the appropriate cell populations. Furthermore, since the IEOs display predominantly vestibular-like features, we identified whether a given gene had been linked to vestibular, as well as auditory phenotypes. We anticipate that the list of target genes present in IEOs will expand as a modified induction platform would allow to yield cochlear supporting cells and hair cells in future studies. Together, our data provide a powerful resource for the community to design and plan SNHL modeling studies using IEOs.

## Discussion

In this study, we differentiate multiple hPSC lines to induce inner ear cell types and used single-cell transcriptomics to evaluate late-stage aggregates of up to 110 days in culture. We also created the first human inner ear atlas at a single-cell level containing data from fetal and adult inner ear tissue. Our data show that differentiation efficiency is largely dependent on the initial BMP-4 concentration and can be determined by epithelial thickness at D3. Using efficiently differentiated aggregates, we discovered the presences of additional off-target cell types (cranial skeletal myocytes, ependymal cells, vascular endothelial cells) and show that most of the cell types have fetal gene expression signatures. Likewise, the on-target formation of hair cells, otic epithelial cells and POM within the IEOs show overlap with the fetal inner ear transcriptomic data. Using the human inner ear atlas, IEOs are composed of vestibular supporting and epithelial cells, and a relative smaller presence of endolymphatic duct/sac cells and cochlear cell types. Interestingly, other nonsensory and cochlear prosensory cells are lacking, indicating that specific inner ear cell types are induced.

The vestibular versus cochlear and sensory versus nonsensory identities are formed early in otic development, with the otic vesicle undergoing these specifications along its patterned axes (Figure 7A-B), reviewed in (Groves and Fekete, 2012; Ohta and Schoenwolf, 2018). Patterning in the dorsoventral axis, leading to separation in vestibular and cochlear fates, is mediated by signals from surrounding tissues: BMP and Wnt versus sonic hedgehog (SHH) (Figure 7A). For patterning in the anterior-posterior axis, however, a gradient of retinoic acid (RA) allows for separation into an anterior domain giving rise to most of the sensory epithelia, and a posterior domain, which gives rise to the sensory epithelia of the posterior crista and mostly nonsensory epithelia. Using this knowledge, the IEOs are mostly composed of vestibular sensory epithelia derived from the dorsal otic vesicle (Figure 7C). Additionally, the induction of vestibulocochlear ganglion-like neurons that form synapses with the hair cells, indicates to the formation of an anterior-ventral part of the otic vesicle (Koundakjian et al., 2007). The formation of a ventral part is in line with the development of a cochlear epithelium within the IEO. This epithelium is restricted to the medial duct floor-like cells and not observed in previous work (Figure 7C) (Koehler et al., 2017). During otic development, the cochlear medial duct floor is mediated by a high activity of Wnt and FGF, with the formation of a prosensory and lateral domain of the duct floor requiring BMP activity (Figure 7B), reviewed in (Munnamalai and Fekete, 2020). Despite sparse literature on prosensory versus nonsensory development in the vestibular organs, BMP activity is not required for proper sensory formation of the cristae and maculae (Ohyama et al., 2010). The presence of Wnt activity and insufficient BMP activity would explain why prosensory development of the cochlea is lacking in the IEOs, but a vestibular sensory domain can exist. The differentiation protocol might benefit from increased BMP activity to induce these prosensory cell types. In addition, increasing SHH activity within the otic vesicle stage of the differentiation protocol might allow for increased cochlear induction within IEOs. Furthermore, the lack of nonsensory epithelia, which are equally important for normal inner ear function, might be overcome by increasing RA activity. Manipulating these pathways might result in more patterned IEOs containing a clinically relevant inner ear cell type diversity.

**Figure 7.**
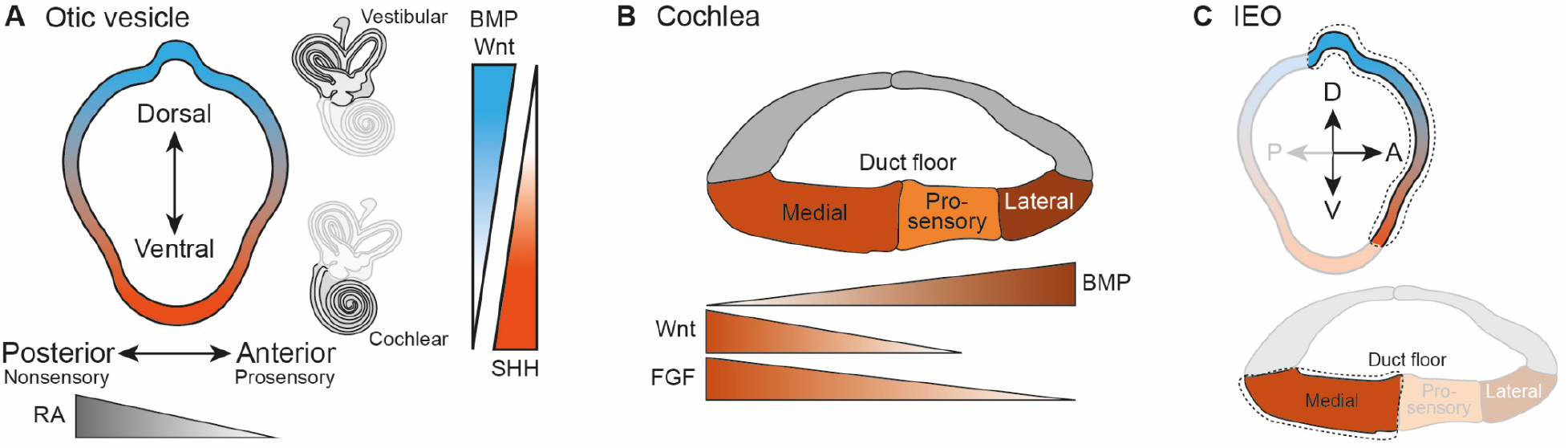
Patterning of the developing otic vesicle and inner ear. **A.** Overview of the signaling pathways that mediate patterning of the otic vesicle in the dorsoventral axis (BMP and Wnt versus SHH) and anterior-posterior axis (gradient of RA). **B.** Signaling pathways which mediate the segregation of the cochlear duct floor into the different domains. **C.** In theory, limited patterning takes place during IEO development, with only specific domains developing (encircled).

Based on the current data, the IEOs can be compared to the human inner ear at approximately W9, with immature type I and type II vestibular hair cells, development of specific cell types of the fetal cochlea and vestibular organs, and presence of melanocytes surrounding the otic epithelia (Van Beelen et al., 2020). We show additional off-target formation of neuroectoderm and surface ectoderm-derived neuroepithelial cells, ependymal cells, keratinocytes, as well as neural crest-derived melanocytes and Schwann cells. Recent work on early IEO culture (<D36) show the formation of these lineages (Steinhart et al., 2021). In this study, mesoderm-derived cranial skeletal myocytes and endothelial cells also develop. One of the challenges of using IEOs in pre-clinical studies is this off-target induction. Off-target chondrocytes for instance, hamper prolonged culture (>D110) of the aggregates as IEOs are lost over time probably due to a decrease in nutritional access and the repressing effect exerted by these off-target cell types. Being able to culture these aggregates for longer time periods could potentially allow for further maturation beyond W9. Alternatively, the use of the endothelial population as potential vasculature together with media flow could enhance maturation as has been described for kidney organoids (Homan et al., 2019).

We present the first human inner ear atlas containing vestibular and cochlear tissue from the first trimester as well as adult vestibular tissue. No single-cell reference of the human inner ear was available, as recent work using the transcriptomics approaches is focused on mouse inner ears (Burns et al., 2015; Jan et al., 2021; Kolla et al., 2020; Korrapati et al., 2019; Wilkerson et al., 2021) or limited to human cochlear tissues (Yu et al., 2019). Future research efforts aim to expand on the human inner ear atlas in order to generate additional resources for the international scientific community in understanding inner ear development and interrogating inner ear development in the context of genetic SNHL. Current knowledge of these pathophysiological mechanisms is often missing, biased towards cochlear neurosensory epithelium, and disregards the spatiotemporal expression of genes and proteins during (human) inner ear development. Additionally, there is increasing awareness for vestibular dysfunction associated with SNHL (Wang et al., 2021). This manuscript motivates the importance of developing human inner ear atlases containing the whole inner ear throughout different stages of development.

Although diversity or maturity of the human inner ear is not fully captured, IEOs could potentially be used for disease modeling. Based on protein and gene expression, a promising SNHL cause to model is for instance congenital Usher syndrome in which the function of hair cells, in both the cochlea and vestibular organs is affected. Within the epithelium also DFNA3A and DFNB1A (*GJB2*) as well as DFNA3B (*GJB6*) can be modeled. Additionally, DFNA9 (*COCH*), DFNA40 (*CRYM*), and DFNX2 (*POU3F4*), all of which affect the function of POM, can potentially be modelled within the IEOs. These are examples of numerous other gene targets that could be evaluated in IEOs. The ability to use the IEOs for disease modeling provides a broadly applicable framework to develop cell or gene therapies for genetic syndromic and nonsyndromic SNHL.

### Limitations of the study

There are several limitations to our study. Firstly, using the scRNAseq and snRNAseq approach we were not able to provide a thorough analysis of cell line differences, nor could we perform a batch-to-batch analysis, since two lines were analyzed by the single-cell method and two lines by the single-nucleus approach. Secondly, given cell type diversity in the IEOs, minor cell populations might not be detected in samples with less than 10,000 cells or nuclei captured per experiment. Thirdly, with merging the single-cell and single-nucleus data, we had to correct for differences in mitochondrial and ribosomal content between methods. Additionally, when analyzing marker gene expression, which is almost entirely based on single-cell references, different expression patters between the single-cell and single-nucleus data are observed. Lastly, due to a shallow single-nucleus sequencing depth, we could not provide a thorough insight into transcription factors as cell identity drivers. Future datasets of both IEO and human fetal and adult specimens, combining the approach of single-cell transcriptomics with spatial biology (e.g., spatial transcriptomics) or chromatin state (e.g., ATAC-sequencing), would allow for more insight into inner ear development. We encourage fellow researchers to use the IEO dataset and human inner ear atlas to make additional discoveries.

## Supporting information

Table S1. Evaluation of 158 SNHL genes and their expression patterns in IEOs.

## Acknowledgements

This work was supported by the Novo Nordisk Foundation (grant NNF21CC0073729), the Hoogenboom-Beckfonds foundation, Erasmus+, ZonMw, and the NIH (grants R01AR075018, R01DC017461, R03DC015624). We thank the Human iPSC Hotel (Leiden University Medical Center, the Netherlands) as well as Dr. M. Sahin and Dr. J. Gonzalez-Heydrich (Boston Children’s Hospital, USA) for sharing their hiPSC lines. We also thank Dr. H. Versnel and F.G.J. Hendriksen (University Medical Center Utrecht, the Netherlands) as well as the Cell Microscopy Core (University Medical Center Utrecht, the Netherlands) for allowing us to use their laboratory facilities. We also acknowledge the Mildred Clinics (Arnhem and Eindhoven, the Netherlands) for supplying inner ear samples, and the Light and Electron Microscopy Facility (Leiden University Medical Center, the Netherlands) for the use of their expertise and microscopes. We also thank T.O. Verhagen, MD and L.C.M. Grijpink, MD (Leiden University Medical Center, the Netherlands) as well as Dr. J. Lee and S.A. Serdy (Boston Children’s Hospital, USA) for their support in preparation of this manuscript. Additionally, we thank Thomas Coate (Georgetown University, USA) for the gift of the POU3F4 antibody. The antibodies developed by Dr. D.J. Orten and Dr. W.E. Wright were obtained from the Developmental Studies Hybridoma Bank, created by the NICHD of the NIH and maintained at The University of Iowa, Department of Biology, Iowa City, IA, USA. Lastly, we acknowledge the Harvard Single Cell Core, the Harvard Chan Bioinformatics Core, the Research Computing Group (Harvard University, USA), and the Leiden Genome Technology Center (Leiden University Medical Center, the Netherlands) for their support and services in obtaining the RNA sequencing data, as well as Dr. S.M. Kielbasa (Leiden University Medical Center, the Netherlands) for SNP demultiplexing of the snRNAseq data.

## Author Contributions

Conceptualization, W.H.V., K.R.K. and H.L.; Methodology, W.H.V, M.R.S., C.N.L., K.R.K. and H.L.; Software, W.H.V. and M.R.S.; Validation, W.H.V., E.S.A.B., M.R.S. and C.N.L.; Formal Analysis, W.H.V.; Investigation, W.H.V., E.S.A.B., M.R.S., C.N.L. and J.C.M.J.G.; Resources, W.H.V., E.S.A.B., M.R.S., C.N.L. and J.C.M.J.G.; Data Curation, W.H.V., K.R.K. and H.L.; Writing – Original Draft, W.H.V., K.R.K. and H.L.; Writing – Review & Editing, W.H.V., E.S.A.B., M.R.S., C.N.L., J.C.M.J.G., P.P.G.B., K.R.K. and H.L.; Visualization, W.H.V. and J.C.M.J.G.; Supervision, P.P.G.B., K.R.K. and H.L.; Project Administration, W.H.V., P.P.G.B., K.R.K. and H.L.; Funding Acquisition, W.H.V., P.P.G.B., K.R.K. and H.L.

## Inclusion and Diversity

We support inclusive, diverse, and equitable conduct of research.

## Material and Methods

### RECOURSE AVAILABILITY

#### Lead contact

Further information and requests for resources and reagents should be directed to and will be fulfilled by the lead contact, Wouter H. van der Valk.

#### Materials availability

This study did not generate new unique reagents.

#### Data and code availability

Single-cell and single-nucleus RNA-seq data are deposited at Gene Expression Omnibus. Accession numbers are GSE214099 and GSE213796, for the IEO data and human inner ear data, respectively. This paper does not report original code. Scripts used for scRNAseq and snRNAseq of IEOs analysis are available at https://github.com/Koehler-Lab/, scripts used for the human inner ear atlas are available at https://github.com/OtoBiologyLeiden/. Any additional information required to reanalyze data reported in this paper is available from the lead contact upon request.

### EXPERIMETNAL MODEL AND SUBJECT DETAILS

#### Pluripotent stem cell lines

Experiments were performed with eight apparently healthy hPSC lines. The WA01 human embryonic stem cell line (male, passages 47-52) was purchased from WiCell Research Institute, provided by Dr. James Thomson at the University of Wisconsin. The two commercially available hiPSC lines that were used are from the same parental male background line (GM25256 from the Coriell Institute): AICS-0074 with the SRY-Box Transcription Factor 2-mEGFP (SOX2-GFP) reporter (hereafter WTC-SOX2, passages 34-46) acquired from the Allen Institute and WTC-GCaMP with a version of the genetically engineered calcium indicator (passages 36-44) provided by Bruce Conklin at the Gladstone Institutes/UCSF (Huebsch et al., 2015). We used additional hiPSC lines that were generated in-house. Generated by the Human iPSC Hotel (Leiden University Medical Center) are LUMC0004iCtrl10 (hereafter LUMC04i10, male, passages 15-36) (Dambrot et al., 2014) and LUMC044iCtrl44 (hereafter LUMC44i44, female, passages 20-43) (Chen et al., 2017). Generated by the Human Neuron Core (Boston Children’s Hospital) are SAH0047-02 (female, passages 23-25) (Ebrahimi-Fakhari et al., 2016), GON0515-03 (male, passages 22-24) (Shlevkov et al., 2019) and GON0926-02 (female, passages 19-21). The cell lines tested negative for mycoplasma prior to experimentation. Cells were cultured on 6-well plates coated with vitronectin recombinant human protein (A14700, Gibco, concentration 0.5 µg/cm) and maintained in Essential 8 Flex medium (A2858501, Gibco) with Normocin (ant-nr-1, Invivogen, concentration 100 µg/ml), hereafter E8 medium. The E8 medium was replenished every other day or every day depending on cell confluency. At approximately 80% confluence (every 4-5 days on average), WTC-SOX2 cells were passaged as single cells using Stempro Accutase Cell Dissociation Reagent (hereafter Accutase; A1110501, Gibco) using 10 µM Y27632 (04-0012-02, Stemgent). The Y27632 was removed within 24 h after passaging by a medium replenishment. All other lines were passaged at approximately 80% confluency (every 4-5 days on average) as tiny clusters (3-5 cells on average) using 0.5 mM EDTA (15575020, Gibco, concentration 10 mM).

#### Human inner ear tissue

Collection of human fetal inner ear tissue was performed according to the Dutch legislation (Fetal Tissue Act, 2001) and the WMA Declaration of Helsinki guidelines. For this project, we obtained approval from the Medical Research Ethics Committee of Leiden University Medical Center (protocol registration number B19.070) as well as written informed consent of the donor following the Guidelines on the Provision of Fetal Tissue set by the Dutch Ministry of Health, Welfare, and Sport (revised version, 2018). The inner ear tissue was collected after elective termination of pregnancy by vacuum aspiration. Embryonic or fetal age was determined by obstetric ultrasonography prior to termination, with a standard error of 2 days.

Adult vestibular tissue was collected during translabyrinthine vestibular schwannoma surgery This project was approved by the Medical Research Ethics Committee of Leiden University Medical Center (protocol registration number B18.028) and tissue was obtained after written informed consent of the donor.

### METHOD DETAILS

#### BMP-4 preparation

BMP-4 from different vendors were used for these experiments. BMP-4 from Peprotech (120-05) was reconstituted in sterile 5 mM citric acid containing 0.2% (wt/vol) of human serum albumin to a concentration of 100 µg/ml. BMP-4 from Stemgent (03-0007) and R&D Systems (314-BPE) were reconstituted in sterile 4 mM HCl to a concentration of 100 µg/ml. BMP-4 from R&D Systems (314-BP) was reconstituted in sterile 4 mM HCl containing 0.1% (wt/vol) human serum albumin to a concentration of 100 µg/ml. Reconstituted BMP-4 was stored as 2 μL aliquots at −80 °C and replaced every six months.

#### Differentiation of hPSCs

Inner ear organoid induction followed the protocol recently published with minor alterations (Koehler *et al*., 2017; Steinhart *et al*., 2021). In brief, colonies of human pluripotent stem cells were detached and dissociated using Accutase and collected as a single-cell suspension in E8 medium containing 20 µM Y27632. 3,500 cells in 100 µl suspension were distributed per well of 96-well U-bottom plates with a super-low cell attachment surface (174925, Thermo Scientific). After centrifugation at 110x*g* for 6 min to aid aggregation, the cell aggregates were incubated in a 37°C incubator under 5% CO_2_ for 48 h. After 24 h, 100 µl of fresh E8 medium was added per well to dilute out Y27632.

At differentiation day 0 (D0), all cell aggregates were collected and individually transferred to a new 96-well U-bottom plate with a super-low cell attachment surface in 100 µl of E6 medium (A1516401, Gibco) with Normocin (concentration 100 µg/ml), hereafter E6 medium, containing 2% Matrigel® Growth Factor Reduced (hereafter Matrigel; 354230, Corning), 10 µM SB431542 (04-0010-05, Stemgent), 4 ng/ml basic FGF (hereafter bFGF; 100-18B, PeproTech) and 0-40 ng/ml BMP-4. On D3, 25 µl of E6 per well was added in with an end-concentration of 200 ng/ml LDN (04-0074-02, Stemgent) and 50 µg/ml bFGF, making the total volume 125 µl per well. This treatment induces cranial neural crest formation. On D6, the total volume is increased to 200 µl by adding 75 µl of fresh E6 medium. On D8, 100 µl of the medium was changed for E6 containing 6 µM CHIR99021 (hereafter CHIR; 04-0004-02, Stemgent) to induce otic placode formation. On D10, 100 µl of the medium was changed for fresh E6 containing 3 µM CHIR. On D12, the aggregates were transferred to Organoid Maturation Medium (OMM) containing 1% Matrigel and 3 µM CHIR in a 24-well plate with a super-low cell attachment surface (174930, Thermo Scientific) on an in-incubator orbital shaker. OMM is a 1:1 mixture of advanced DMEM:F12 (12634010, Gibco) and Neurobasal Medium (21103049, Gibco), supplemented with 0.5x B27 without vitamin A (12587010, Gibco), 0.5x N2 Supplement (17502048, Gibco), 1x Glutamax (35050061, Gibco), 0.1 mM ß-mercaptoethanol (21985023, Gibco), and Normocin. On D15, a half medium change was performed with OMM containing 1% Matrigel and 3 µM CHIR. On D18, a half medium change with OMM was performed, diluting out the Matrigel and CHIR. On D21, a full medium change with OMM was performed. Half of the medium was replenished every 3 days, with a full medium change every 9 days. The volume was gradually increased starting from 500 µl at D12, to 1 ml on approximately D45, to 1.5 ml on approximately D60 and later.

#### *In vitro* measurements of aggregate morphology

Aggregates were monitored during the differentiation period with a Nikon TS2R microscope or an EVOS M5000 microscope using either bright-field or phase-contrast visualization. Images collected were analyzed using ImageJ measuring epithelial thickness at three different areas of the aggregate, circularity and area.

#### Vibratome-slicing of aggregates

Aggregates were embedded in low 2% low melting point agarose (16520050, Invitrogen) and 200 µm thick slices were obtained using a motorized vibratome (VT1200, Leica Microsystems). Sections were obtained in ice-cold PBS (14190144, Gibco) and bright-field evaluation was performed using a Nikon TS2R microscope. Selected sections were processed for scRNAseq as described below.

#### Aggregate dissociation to single cells for scRNAseq

Vibratome sections of randomly selected aggregates were pooled at specified time points. Three to four number of aggregates were placed in warm TrypLE solution (12605036, Gibco) and incubated in a 37°C incubator under 5% CO_2_ on an orbital shaker set to 65 rpm. The mixture was triturated with a pipette using wide bore p1000 tips every 10 min. Dissociation to single cells was complete after 60 min and confirmed by bright-field visualization using a Nikon TS2R. TrypLE activity was stopped by adding cold 3% bovine serum albumin (BSA; A7030, Sigma-Aldrich) in PBS. To remove residual cell aggregates and debris, the suspension was filtered with a 40 μm Flowmi cell strainer (H13680-0040, Bel-Art). After washing three times with 3% BSA in PBS, cells were resuspended in ice-cold 3% BSA in PBS and filtered again with a 40 μm Flowmi cell strainer. Cell viability was determined by using Trypan Blue (15250061, Gibco) and an automated cell counter (Countess II, Thermo Fisher). Cell suspensions were diluted with cold 3% BSA in PBS to reach a final concentration of 1,000 cells/μl.

#### Aggregate dissociation to single nuclei for snRNAseq

Two to four randomly selected aggregates were processed at specified time points. Aggregates were pooled, washed once with ice-cold PBS, and resuspended in lysis buffer that contained 10 mM Tris-HCl (T2194, Millipore-Sigma), 10 mM NaCl (59222C, Millipore-Sigma), 3 mM MgCl_2_ (M1028, Millipore-Sigma), 0.1% Nonidet P40 Substitute (74385, Millipore-Sigma) in DEPC-treated water (750024, Invitrogen). The suspension was transferred to a Dounce tissue grinder (885300-0002, Kimble) and ground every 5 min. After 20 min, dissociation to single nuclei was confirmed by bright-field visualization (EVOS M5000, Thermo Scientific). After trituration using p1000 and p200 tips, the suspension was filtered through a 70 μm MACS SmartStrainer (130-098-462, Miltenyi Biotec), centrifuged at 500x*g* for 5 min at 4°C and resuspended in 1% BSA in PBS (AM2616, Invitrogen) containing 200 U/μl RNase inhibitor (3335402001, Millipore-Sigma). After filtration using a 40 μm Flowmi cell strainer and centrifugation at 500g for 5 min at 4°C, the nuclei suspension was resuspended to a final concentration of 1,000 cells/μL.

For one experiment of D110 aggregates generated using the LUMC04i10 hiPSC line, we performed nuclear hashtagging of two separate aggregates using single nuclei hashing antibodies (MAb414, Merck). To this end, the nuclei suspension was blocked using Fc-blocking reagent (422302, Biolegend) for 5 min on ice followed by incubation with the hashing antibody (1 µg per 100 µl) for 10 min on ice. Subsequently, nuclei were washed three times using 1% BSA in PBS containing 200 U/μl RNase inhibitor, pooled together 1:1, and resuspended to a final concentration of 1,000 cells/μl.

#### Human inner ear tissue dissociation for snRNAseq

Human fetal inner ear tissue was stored in RNAlater (AM7020, Invitrogen) at 4°C and processed to a single nuclei suspension within 16 h after collection. Human adult tissue was processed within 20 min after collection. After collection of the inner ear tissue, the tissue was evaluated using a dissection microscope (M205C, Leica). The otic capsule of the W9.2 fetal inner ear was removed. The human inner ear tissue followed a similar protocol of the aggregate dissociation to single nuclei as described above. Minor alterations include the lysis buffer that contains 0.005% Nonidet P40 substitute and additional filtration steps following the 40 μm Flowmi cell strainer: the suspension was filtered through a 20 μm and 10 μm Pluristrainer (43-10020-40 and 43-10010-40, PluriSelect) to remove debris.

#### RNA-seq cDNA library preparation and sequencing

Single cell and nucleus gene expression libraries were generated on the 10x Genomics Chromium platform using the Chromium Next GEM Single Cell 3’ Library & Gel Bead Kit v2 (scRNAseq) or v3.1 (snRNAseq) and Chromium Next GEM Chip G Single Cell Kit (10x Genomics) according to the manufacturer’s protocol. Hashtag libraries were amplified and barcoded as in Stoeckius et al. (Stoeckius et al., 2018). Gene expression and hashtag libraries were sequenced on a NextSeq500 or NovaSeq 6000 S4 flow cell using v1 Chemistry (Illumina) and FASTQ files were generated using Cell Ranger mkfastq (10x Genomics).

#### Analysis of scRNAseq and snRNAseq data

The 10x Genomics Cell Ranger 6.1.0 pipeline (http://support.10xgenomics.com/) was used to generate demultiplexed FASTQ files from the raw sequencing reads. The reads were aligned to the human reference GRCh38 genome. For snRNAseq reads, reads were mapped to introns as wells as exons, resulting in a greater number of genes detected per nucleus. Levels of gene expression were analyzed using the number of UMIs (unique molecular indices) detected in each cell. Cell Ranger was used to generate filtered gene expression matrices.

Low-quality cells were removed by quality control on UMI number, number of genes, and percentage of transcripts aligned to mitochondrial genes as well as ribosomal genes. The parameters used were different between the single-cell and single-nucleus data. After normalization and scaling of the data, identification of variable features was performed using SCTransform (Hafemeister and Satija, 2019) regressing out the effect of the effect of cell cycle, mitochondrial content and/or ribosomal content.

Following principal component analysis (PCA) determining the most highly variable genes, Harmony was used to integrate datasets (Korsunsky et al., 2019). These integrated datasets were used to visualize clustering by Uniform Manifold Approximation and Projection (UMAP) plotting techniques. Cell cluster identification was performed using differential expression analysis and manually by using cell type-specific marker genes.

R packages ggplot2 and Nebulosa were used to plot quantitative gene expression and specific gene expression patterns. Symphony was used to map the integrated IEO scRNAseq and snRNAseq data to the human inner ear atlas (Kang *et al*., 2021).

#### Aggregate processing and immunohistochemistry

Samples were washed twice with PBS and fixed using 4% formaldehyde (1.04005.1000, Merck) in 0.1 M Na^+^/K^+^-phosphate buffer (pH 7.4) overnight at 4°C. Fixed samples were either paraffin embedded or processed for cryosectioning. For paraffin embedding, aggregates were dehydrated in an ascending ethanol (70%–99%; 84050068.2500, Boom) series, cleared in xylene (534056, Honeywell), and embedded in paraffin wax (2079, Klinipath). Sections of 4-5 µm were cut using a rotary microtome (HM355S, Thermo Scientific). Sections were deparaffinized in xylene and rehydrated in a descending series of ethanol (96%–50%) and several rinses in deionized water. At approximately every 10 sections, one section was selected for routine staining with hematoxylin (40859001, Klinipath) and eosin (40829002, Klinipath). Before immunostaining, antigen unmasking was performed either in 10 mM sodium citrate buffer (pH 6.0; S1804-500G, Sigma-Aldrich) for 12 min at 97°C. For cryosectioning, aggregates were cryoprotected with graded treatments of 15% and 30% sucrose solutions in PBS and embedded in tissue-freezing medium (1518313, General Data Healthcare) using cryomolds (25608, Andwin Scientific). Cryosections of 12 µm were cut using a cryostat (CM-1860, Leica Microsystems). For immunostaining, sections were rinsed in washing buffer consisting of 0.05% Tween-20 (H5152, Promega) and incubated for 30 min with blocking solution consisting of 5% bovine serum albumin (BSA; A7030, Sigma-Aldrich) and 0.05% Tween-20 in PBS, followed by overnight incubation at 4°C with the primary antibodies listed below. Next, sections were incubated at ambient temperature (RT) with the Alexa Fluor-conjugated secondary antibodies for 2 h, listed below. Nuclei were stained with 4′,6-diamidino-2-phenylindole (DAPI; 1:1,000; D1306, Invitrogen). Sections were mounted with ProLong™ Gold Antifade Mountant (P36934, Invitrogen). Negative controls were carried out by matching isotype controls and/or omitting primary antibodies. Positive controls were carried out by staining sections of known positive human tissue samples based upon their protein expression profiles, including human inner ear tissue. At least two separate immunostaining experiments were performed with each primary antibody.

| Antibody | Source | Identifier |
| --- | --- | --- |
| Primary |  |  |
| Mouse monoclonal anti-ACTC1 (1:50) | R&D systems | Cat#: MAB9308-100 |
| Mouse monoclonal anti-ANXA4 (1:200) | Sigma-Aldrich | Cat#: SAB4200121 |
| Mouse monoclonal anti-ATP1A1 (1:200) | Novus Biologicals | Cat#: NB300-146 |
| Rabbit polyclonal anti-BSND (1:750) | Novus Biologicals | Cat#: NBP2-49101 |
| Mouse monoclonal anti-CDH1 (1:50) | BD Transduction Laboratories | Cat#: 610182 |
| Rabbit polyclonal anti-CDH23 (1:50) | Invitrogen | Cat#: PA5-53564 |
| Mouse monoclonal anti WRHN (1:50) | Abnova | Cat#: H00025861-M05 |
| Rabbit polyclonal anti-GJB2 (1:100) | Alomone Labs | Cat#: ACC-212 |
| Rabbit polyclonal anti-GJB6 (1:100) | Invitrogen | Cat#: PA5-11640 |
| Mouse monoclonal anti-MLANA (1:50) | Abcam | Cat#: ab731 |
| Rabbit polyclonal anti-MYO7A (1:100) | Proteus Biosciences | Cat#: 25-6790 |
| Rabbit monoclonal anti-MYO7A (1:50) | Novus Biologicals | Cat#: NBP1-84266 |
| Mouse monoclonal anti-MYO7A (1:30) | DSHB | Cat#: 138-1s |
| Mouse monoclonal anti-MYOG (1:10) | DSHB | Cat#: F5D |
| Rabbit polyclonal anti-OCM (1:200) | Abcam | Cat#: ab150947 |
| Rabbit polyclonal anti-OTOF (1:100) | Novus Biologicals | Cat#: NBP1-46574 |
| Rabbit polyclonal anti-OTOR (1:100) | Invitrogen | Cat#: PA5-55053 |
| Goat polyclonal anti-PDGFRB (1:15) | R&D systems | Cat#: AF385 |
| Mouse monoclonal anti-PECAM1 (1:50) | Dako | Cat#: M082329-2 |
| Alexa Fluor™ 568 phalloidin | Invitrogen | Cat#: A12380 |
| Chicken ...clonal anti-POU3F4 (1:10000) | Aves Labs | Cat#: AB_2814704 |
| Rabbit polyclonal anti-SIX1 (1:100) | Invitrogen | Cat#: PA5-51654 |
| Rabbit polyclonal anti-SLC26A4 (1:100) | Novus Biologicals | Cat#: NBP1-85237 |
| Goat polyclonal anti-SOX10 (1:50) | Invitrogen | Cat#: PA5-47001 |
| Mouse monoclonal anti-SOX2 (1:100) | BD Transduction Laboratories | Cat#: 561469 |
| Rabbit monoclonal anti-TFAP2A (1:50) | Abcam | Cat#: ab108311 |
| Rabbit polyclonal anti-TNNT1 (1:500) | R&D systems | Cat#: NBP2-38855 |
| Rabbit monoclonal anti-TTR (1:50) | Invitrogen | Cat#: MA5-32634 |
| Mouse monoclonal anti-TUBB3A (1:100) | Biolegend | Cat#: 801202 |
| Mouse monoclonal anti-TUBB3A (1:100) | R&D Systems | Cat#: MAB1195 |
| Secondary |  |  |
| Donkey anti-mouse Alexa Fluor™ 488 (1:1000-2000) | Invitrogen | Cat#: A21202 |
| Donkey anti-mouse Alexa Fluor™ 568 (1:1000-2000) | Invitrogen | Cat#: A10037 |
| Donkey anti-mouse Alexa Fluor™ 594 (1:1000-2000) | Invitrogen | Cat#: A21203 |
| Donkey anti-mouse Alexa Fluor™ 647 (1:1000-2000) | Invitrogen | Cat#: A31571 |
| Donkey anti-mouse Alexa Fluor™ 680 (1:1000-2000) | Invitrogen | Cat#: A10038 |
| Goat anti-mouse IgG1 Alexa Fluor™ 488 (1:2000) | Invitrogen | Cat#: A21121 |
| Goat anti-mouse IgG1 Alexa Fluor™ 568 (1:2000) | Invitrogen | Cat#: A21124 |
| Goat anti-mouse IgG1 Alexa Fluor™ 647 (1:2000) | Invitrogen | Cat#: A21240 |
| Goat anti-mouse IgG2a Alexa Fluor™ 488 (1:2000) | Invitrogen | Cat#: A21131 |
| Goat anti-mouse IgG2a Alexa Fluor™ 568 (1:2000) | Invitrogen | Cat#: A21134 |
| Goat anti-mouse IgG2a Alexa Fluor™ 647 (1:2000) | Invitrogen | Cat#: A21241 |
| Goat anti-mouse IgG2b Alexa Fluor™ 488 (1:2000) | Invitrogen | Cat#: A21141 |
| Goat anti-mouse IgG2b Alexa Fluor™ 568 (1:2000) | Invitrogen | Cat#: A21144 |
| Goat anti-mouse IgG2b Alexa Fluor™ 647 (1:2000) | Invitrogen | Cat#: A21242 |
| Donkey anti-rabbit Alexa Fluor™ 488 (1:1000) | Abcam | Cat#: ab150061 |
| Donkey anti-rabbit Alexa Fluor™ 488 (1:1000-2000) | Invitrogen | Cat#: A21206 |
| Donkey anti-rabbit Alexa Fluor™ 568 (1:1000-2000) | Invitrogen | Cat#: A10042 |
| Donkey anti-rabbit Alexa Fluor™ 594 (1:1000-2000) | Invitrogen | Cat#: A21207 |
| Donkey anti-rabbit Alexa Fluor™ 647 (1:1000-2000) | Invitrogen | Cat#: A31573 |
| Donkey anti-rabbit Alexa Fluor™ 680 (1:1000-2000) | Invitrogen | Cat#: A10043 |
| Donkey anti-goat Alexa Fluor™ 488 (1:1000) | Abcam | Cat#: ab150133 |
| Goat anti-chicken Alexa Fluor™ 594 (1:1000) | Invitrogen | Cat#: A11042 |

#### Sample processing for transmission electron microscopy

At D74, LUMC04i10 IEOs (n=6) were collected and fixed overnight with 2.5% glutaraldehyde in 0.1 M sodium cacodylate buffer (pH 7.4) at 4°C. Fixed specimens were rinsed (3 x 20 min) in 0.1 M sodium cacodylate buffer (pH 7.4) at RT prior to post-fixation in a solution containing 1% OsO_4_ and 1% K_4_Ru(CN)_6_ (potassium hexacyanoruthenate (II) hydrate) in 0.1 M sodium cacodylate buffer (pH 7.4) for 2 h at 4°C followed by several washes in distilled water. Dehydration was performed in an ascending ethanol series (70%-100) and 100% 1,2-propylene oxide followed by infiltration with a mixture of Spurr’s low-viscosity resin and 1,2-propylene oxide (1:1) for 1 h and fresh pure resin under continuous agitation for 2 h at RT. Finally, each specimen was separately embedded in fresh pure resin in a polyethylene embedding mold and polymerized overnight at 70°C.

#### Light microscopy

For light-microscopical assessment, semi-thin (1 μm) plastic sections were cut with a diamond knife (Histo, DiATOME) on a microtome (RM2265, Leica Microsystems), collected on aminoalkylsilane-coated glass slides, stained with a warm aqueous solution containing 1% methylene blue, 1% azure II and 1% sodium tetraborate, and mounted in Entellan® (1079610100, Merck) under a glass coverslip.

#### Image acquisition and processing

Sections stained with either HE or methylene blue-azure II were examined with a Leica DM5500B microscope equipped with a Leica DFC450C digital color camera. Digital images were acquired and stored using Leica Application Suite software (LAS version 4.5; Leica Microsystems GmbH). Images were adjusted for optimal contrast and brightness and assembled into figures using Adobe Illustrator 2021 or Adobe Photoshop 2021 (Adobe Systems Software Ltd).

Immunostained sections were acquired with a Leica SP8 confocal laser scanning microscope using Leica objectives (20×/0.7 dry HC PL Apo, 40×/1.3 oil HC PL Apo CS2, 63×/1.4 oil HC PL Apo or 100×/1.3 oil HC PL Fluotar), operating under Leica Application Suite X microscope software (LAS X, Leica Microsystems). Maximal projections were obtained from image stacks with optimized z-step size. Brightness and contrast adjustments were performed with Fiji (ImageJ version 1.52p).

#### Transmission electron microscopy

For ultrastructural examination, representative or interesting areas were selected based on light-microscopical assessment. Ultrathin (60-90 nm) sections were cut with a diamond knife (Ultra 45°, DiATOME) on an Ultrotome III (LKB) and collected upon Pioloform^®^-coated, single-slot copper grids. The sections were contrast-stained with 7% uranyl acetate in 70% methanol and Reynolds’ lead citrate solution and examined in a FEI Tecnai™ 12 transmission electron microscope (operating at 80 kV) equipped with a Veleta 4-megapixel side-mount CCD camera (Olympus Soft Imaging Solutions GmbH). Digital images were acquired with TIA software (TEM Imaging & Analysis, version 4.7 SP3; FEI). Alternatively, sets of serial digital images with 10% overlap (at an original magnification of x30,000) were acquired with the automated EM data acquisition software package SerialEM (https://bio3d.colorado.edu/SerialEM). Final alignment (stitching) of the sets of overlapping images and montage blending were done using the reconstruction and modelling software package IMOD (https://bio3d.colorado.edu/imod) operating under the Unix toolkit Cygwin. Montages or single images were finally saved as TIFF and imported into Fiji ImageJ for further image processing, examination and selection of interesting areas. Images were assembled into figures using Adobe Photoshop.

## QUANTIFICATION AND STATISTICAL ANALYSIS

For analyzing the *in vitro* measurements of aggregate morphology, statistical significance was analyzed with 2-way ANOVA with a Sidak correction for multiple comparisons. Data was considered statistically significant if P<0.05.

---

**Figure S1.**
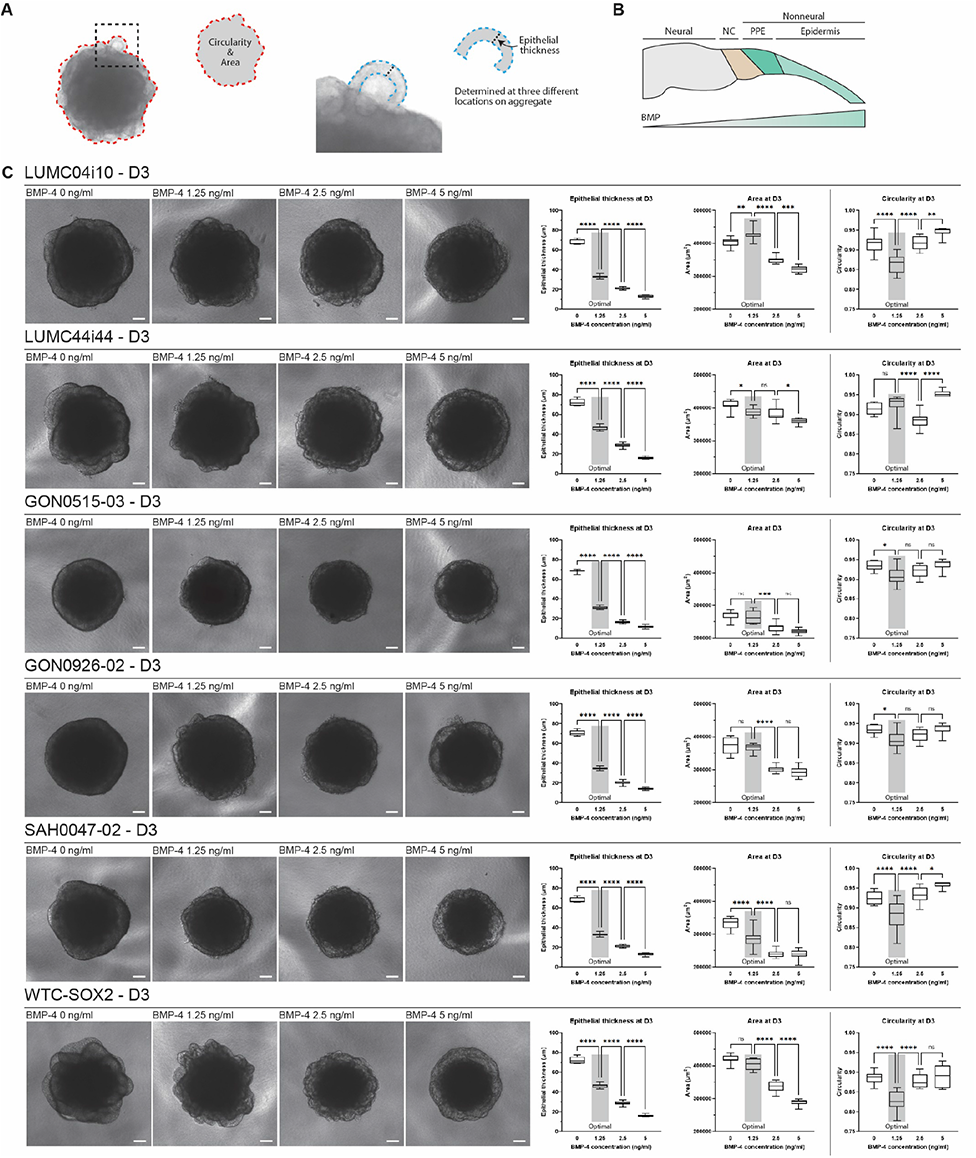
Analyses of differentiation efficiency using morphological characteristics of early IEO-differentiation. **A.** Measurements of D3 aggregates using ImageJ: circularity, area and epithelial thickness. Epithelial thickness was measured at three different locations in the aggregate. **B.** Illustration showing relative thickness difference during ectodermal specification at the gastrula stage which is achieved by a BMP-4 gradient. **C.** Representative phase-contrast images of D3 aggregates treated with different concentrations of BMP-4 for all hiPSC lines. Results of circularity, area and epithelial thickness are plotted per cell line. Statistical significance was analyzed with 2-way ANOVA with a Sidak correction for multiple comparisons. Data was considered statistically significant if P<0.05. n=10 per datapoint was graphed in a min-max box plot. A representative graph of at least 2 individual experiments is shown. The optimal concentration for this set of experiments is highlighted. Scale bars 100 μm. NC: neural crest; PPE: pre-placodal ectoderm.

**Figure S2.**
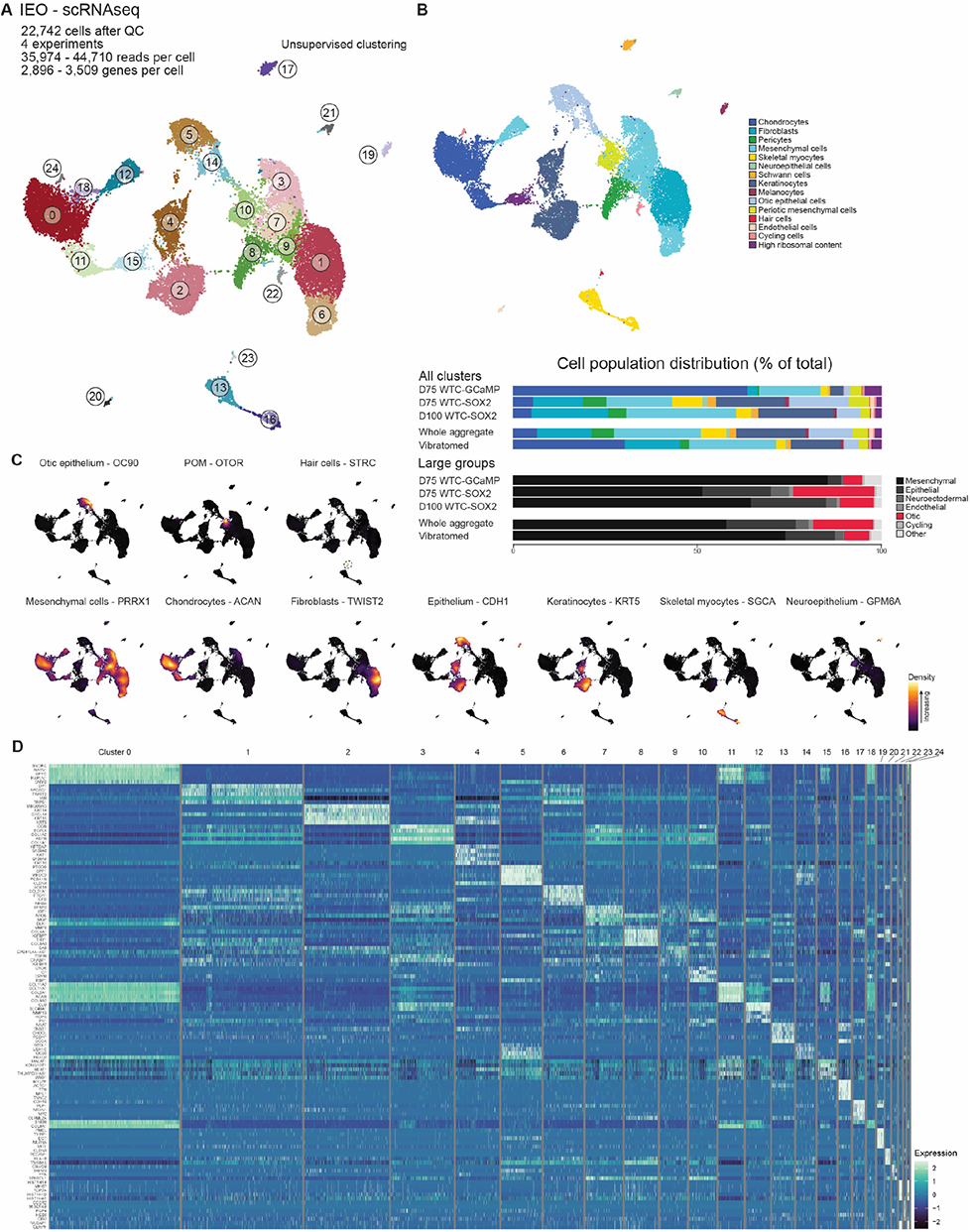
scRNAseq unravels the cell type diversity of D75-D100 IEOs. **A.** Overview UMAP plot of D75 (WTC-GCaMP and WTC-SOX2) and D100 (WTC-SOX2) dataset of 22,742 cells. **B.** Cell type annotation with relative cell population contribution of cell types and large groups. **C.** Marker genes involved in cell type annotation assignment. **D.** Heatmap showing expression patterns of the top 5 differentially expressed genes per unsupervised cluster.

**Figure S3.**
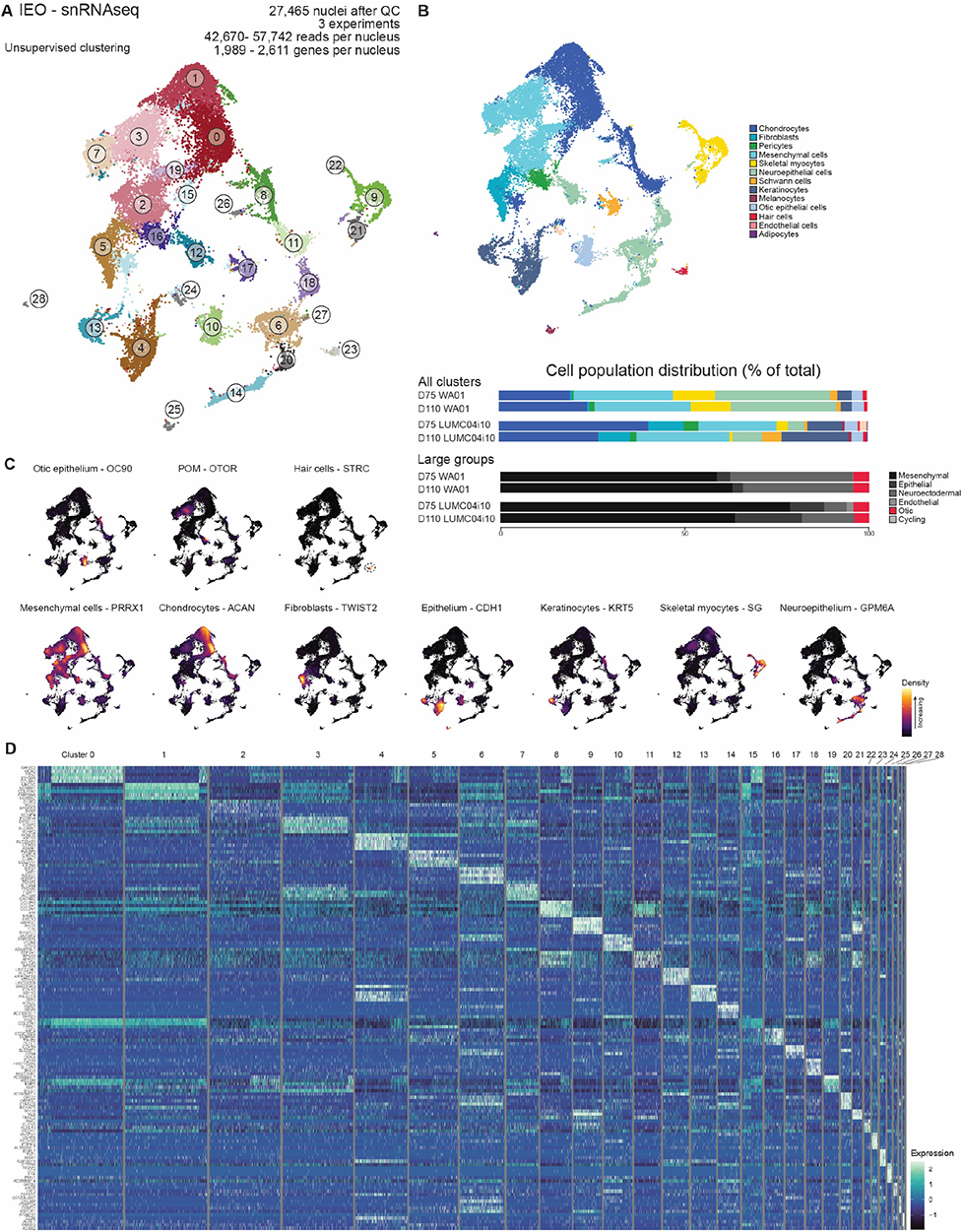
snRNAseq of D75-D110 aggregates shows a similar cell type diversity. **A.** Overview UMAP plot of D110 (WA01, LUMC04i10) dataset of 27,465 nuclei. **B.** Cell type annotation with relative cell population contribution of cell types and large groups. **C.** Marker genes involved in cell type annotation assignment. **D.** Heatmap showing expression patterns of the top 5 differentially expressed genes per unsupervised cluster.

**Figure S4.**
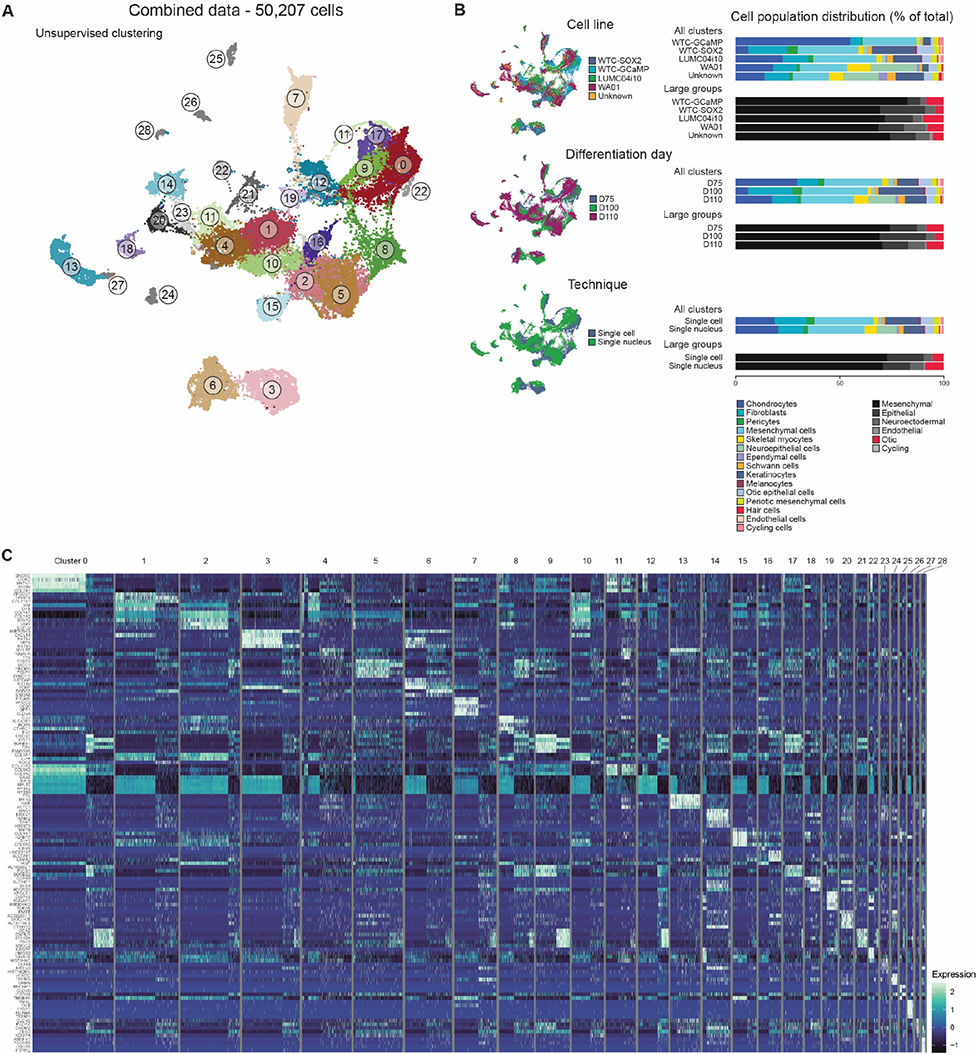
Integration of the scRNAseq and snRNAseq IEO datasets. **A.** Overview UMAP plot of the integrated combined dataset composed of 50,207 cells and nuclei of D75 (WTC-GCaMP, WTC-SOX2, WA01, LUMC04i10), D100 (WTC-SOX2) and D110 (WA01 and LUMC04i10) aggregates. **B.** UMAP plots and relative cell population contribution showing overlap between the cell lines, differentiation days and techniques used. Cell line “Unknown” means that using SNP demultiplexing, no distinction could be made between LUMC04i10 or WA01. **D.** Heatmap showing expression patterns of the top 5 differentially expressed genes per unsupervised cluster.

**Figure S5.**
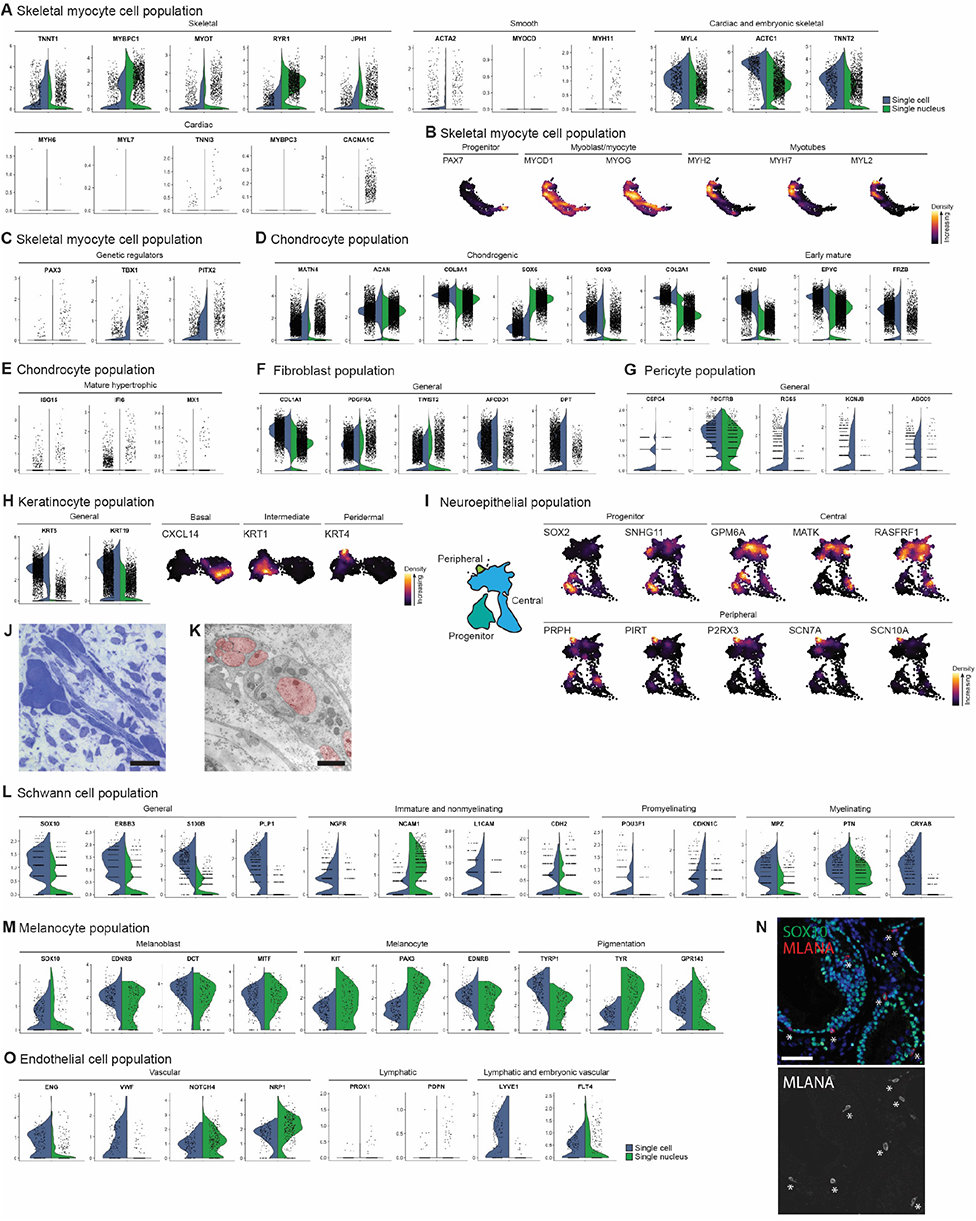
Cell type and development-specific marker gene expression of cell types in D75-D110 aggregates. **A.** Marker gene expression within the skeletal myocyte population of skeletal myocytes, smooth muscle myocytes, cardiomyocytes, and of both cardiomyocytes and fetal skeletal myocytes. **B.** Density plots of the skeletal myocyte cell cluster, showing expression of progenitor, myoblast and myotube markers. **C.** Gene expression within the skeletal myocyte cell population of genetic regulators of caudal (*PAX3*) and cranial skeletal myocytes (*TBX1*, *PITX2*). **D.** Marker gene expression within the chondrocyte population consisting of chondrogenic and early mature chondrocytes. **E.** Marker gene expression within the chondrocyte population of mature hypertrophic markers. **F.** Marker gene expression within the fibroblast population. **G.** Marker gene expression within the pericyte population. **H.** Marker gene expression within the keratinocyte population and density plots showing expression of basal, intermediate and peridermal markers. **I.** Schematic dividing the neuroepithelial cluster into progenitor, central and peripheral neuroepithelial cells based on density plots of the neuroepithelial cell cluster, showing expression of progenitor, central and peripheral identities. **J.** Staining (methylene blue-azure II) depicting a group of neurons surrounded by glial (Schwann) cells lying freely in the mesenchymal stroma in close proximity to an IEO-vesicle within a D74 aggregate. Scale bar 25 μm. **K.** TEM image showing neurons (in red) surrounded by a Schwann cell situated underneath IEO-vesicle within a D74 aggregate. Scale bar 1 μm. **L.** Marker gene expression within the Schwann cell population of immature, nonmyelinating, promyelinating and myelinating markers. **M.** Marker gene expression within the melanocyte population containing melanoblast, melanocyte and pigmented stages. **N.** MLANA^+^ melanocytes in proximity of SOX10^+^ IEO-vesicles in a D75 aggregate. Asterisks show location of melanocytes. Scale bar 50 μm. **O.** Marker gene expression within the endothelial cell population of vascular, lymphatic and combined lymphatic and fetal vascular genes.

**Figure S6.**
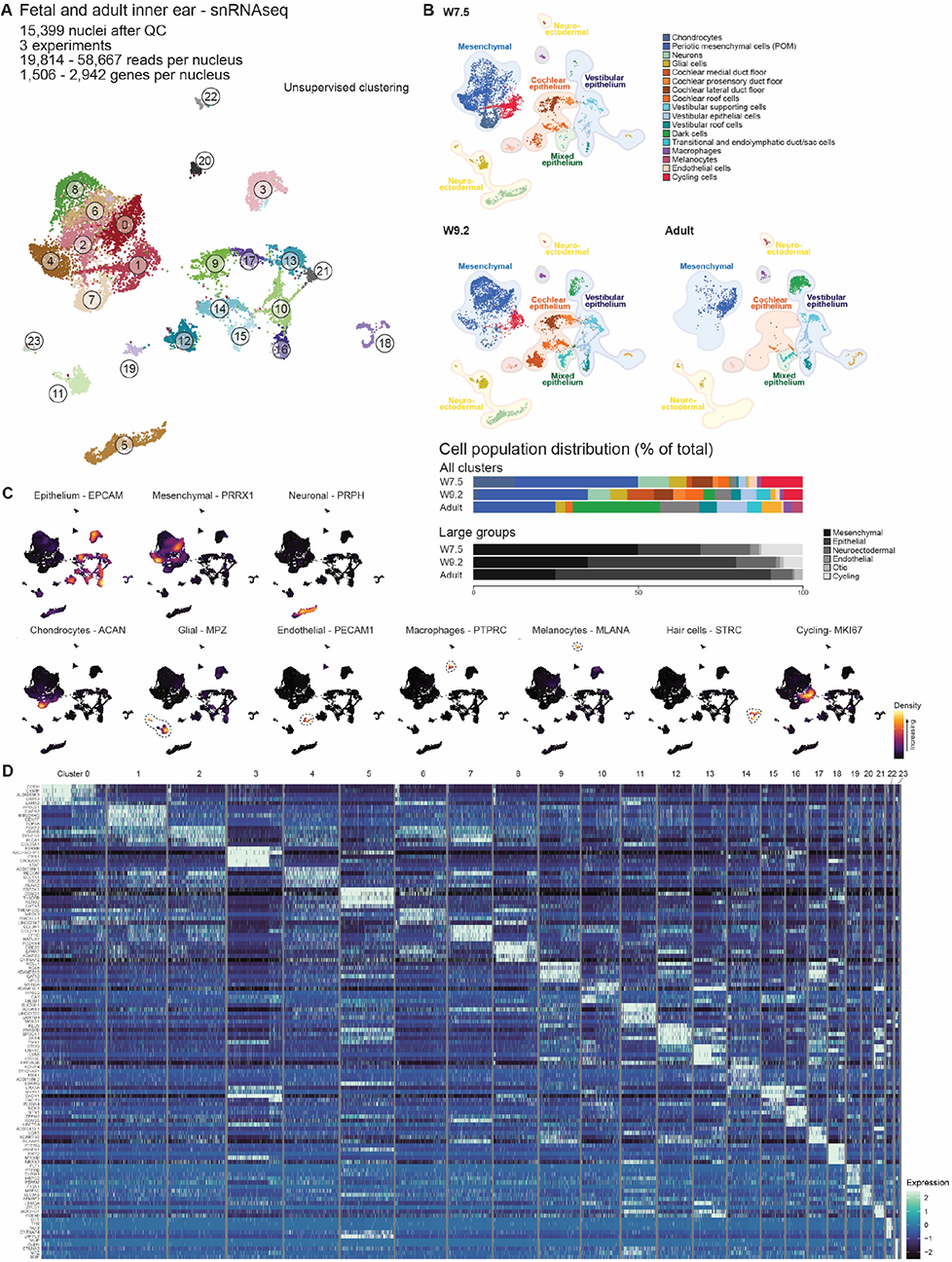
snRNAseq of fetal and adult human inner ear tissue. **A.** Overview UMAP plot of the fetal stages (W7.5, W9.2) and adult inner ear tissue combined dataset of 15,399 cells. **B.** Cell type annotated UMAP plots and relative cell population contribution per developmental timepoint. **C.** Marker genes involved in cell type annotation assignment. **D.** Heatmap showing expression patterns of the top 5 differentially expressed genes per unsupervised cluster.

**Figure S7.**
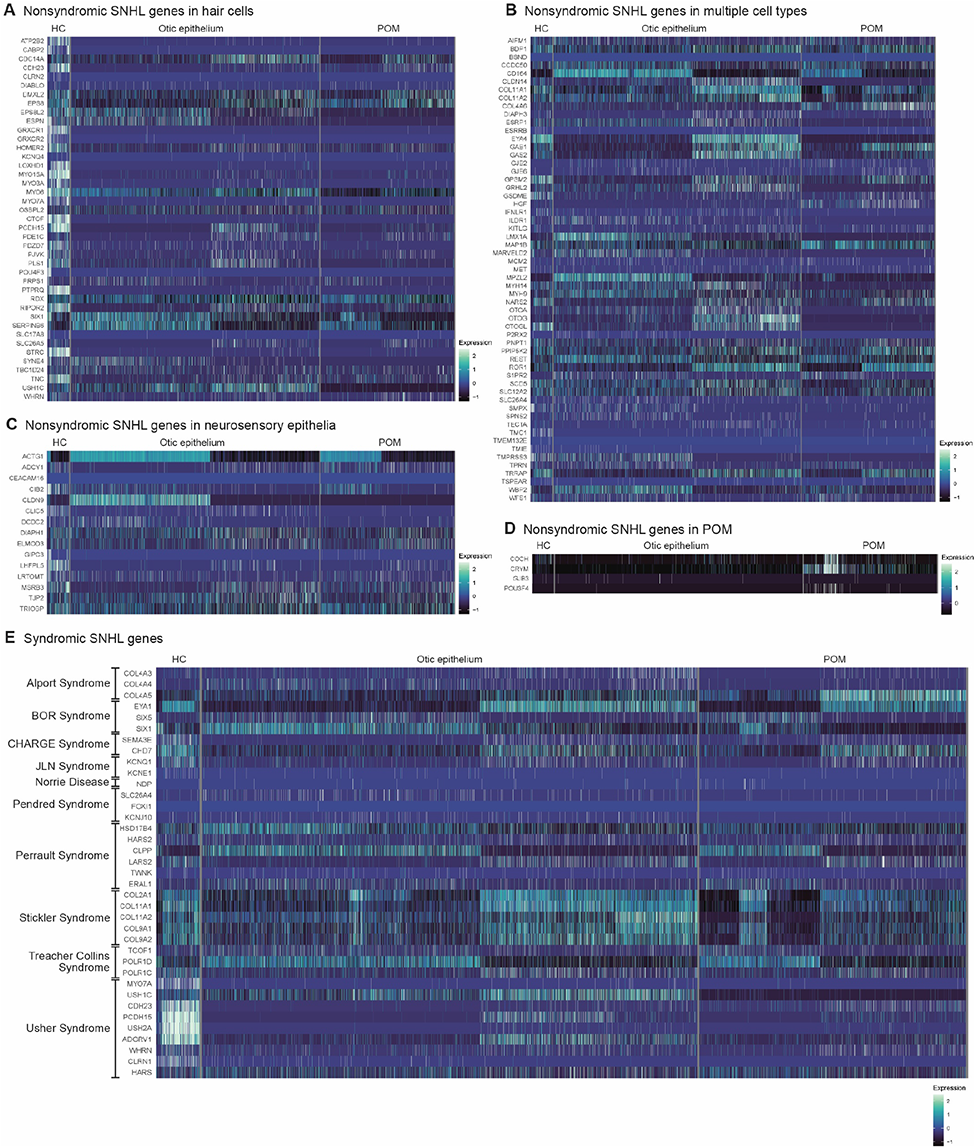
Gene expression of SNHL genes in the otic populations of the D75-D110 IEOs. **A.** Heatmap showing expression patterns of nonsyndromic SNHL genes expressed in hair cells. **B.** Heatmap showing expression of nonsyndromic SNHL genes expressed in multiple inner ear cell types. **C.** Heatmap showing expression of nonsyndromic SNHL genes expressed in neurosensory epithelia. **D.** Heatmap showing expression of nonsyndromic SNHL genes expressed in POM. **E.** Heatmap showing expression of a selection of syndromic SNHL genes. BOR: Branchio-Oto-Renal syndrome; HC: hair cell; JLN: Jervell & Lange-Nielsen syndrome; POM: periotic mesenchyme.

## References

Anderson, P.A., Malouf, N.N., Oakeley, A.E., Pagani, E.D., and Allen, P.D. (1991). Troponin T isoform expression in humans. A comparison among normal and failing adult heart, fetal heart, and adult and fetal skeletal muscle. Circ Res 69, 1226–1233. 10.1161/01.res.69.5.1226.

Bakken, T.E., Hodge, R.D., Miller, J.A., Yao, Z., Nguyen, T.N., Aevermann, B., Barkan, E., Bertagnolli, D., Casper, T., Dee, N., et al. (2018). Single-nucleus and single-cell transcriptomes compared in matched cortical cell types. PLoS One 13, e0209648. 10.1371/journal.pone.0209648.

Belote, R.L., Le, D., Maynard, A., Lang, U.E., Sinclair, A., Lohman, B.K., Planells-Palop, V., Baskin, L., Tward, A.D., Darmanis, S., and Judson-Torres, R.L. (2021). Human melanocyte development and melanoma dedifferentiation at single-cell resolution. Nat Cell Biol 23, 1035–1047. 10.1038/s41556-021-00740-8.

Berendam, S.J., Koeppel, A.F., Godfrey, N.R., Rouhani, S.J., Woods, A.N., Rodriguez, A.B., Peske, J.D., Cummings, K.L., Turner, S.D., and Engelhard, V.H. (2019). Comparative transcriptomic analysis identifies a range of immunologically related functional elaborations of lymph node associated lymphatic and blood endothelial cells. Front Immunol 10, 816. 10.3389/fimmu.2019.00816.

Burns, J.C., Kelly, M.C., Hoa, M., Morell, R.J., and Kelley, M.W. (2015). Single-cell RNA-Seq resolves cellular complexity in sensory organs from the neonatal inner ear. Nature Communications 6, 8557. 10.1038/ncomms9557.

Butcher, E., Dezateux, C., Cortina-Borja, M., and Knowles, R.L. (2019). Prevalence of permanent childhood hearing loss detected at the universal newborn hearing screen: Systematic review and meta-analysis. PLoS One 14, e0219600. 10.1371/journal.pone.0219600.

Chai, R., Xia, A., Wang, T., Jan, T.A., Hayashi, T., Bermingham-McDonogh, O., and Cheng, A.G. (2011). Dynamic expression of Lgr5, a Wnt target gene, in the developing and mature mouse cochlea. J Assoc Res Otolaryngol 12, 455–469. 10.1007/s10162-011-0267-2.

Chen, X., Janssen, J.M., Liu, J., Maggio, I., ‘T Jong, A.E.J., Mikkers, H.M.M., and Gonçalves, M. (2017). In trans paired nicking triggers seamless genome editing without double-stranded DNA cutting. Nat Commun 8, 657. 10.1038/s41467-017-00687-1.

Choi, I.Y., Lim, H., Cho, H.J., Oh, Y., Chou, B.K., Bai, H., Cheng, L., Kim, Y.J., Hyun, S., Kim, H., et al. (2020). Transcriptional landscape of myogenesis from human pluripotent stem cells reveals a key role of TWIST1 in maintenance of skeletal muscle progenitors. Elife 9. 10.7554/eLife.46981.

Czajkowski, A., Mounier, A., Delacroix, L., and Malgrange, B. (2019). Pluripotent stem cell-derived cochlear cells: A challenge in constant progress. Cell Mol Life Sci 76, 627–635. 10.1007/s00018-018-2950-5.

Dambrot, C., Braam, S.R., Tertoolen, L.G., Birket, M., Atsma, D.E., and Mummery, C.L. (2014). Serum supplemented culture medium masks hypertrophic phenotypes in human pluripotent stem cell derived cardiomyocytes. J Cell Mol Med 18, 1509–1518. 10.1111/jcmm.12356.

Delmaghani, S., and El-Amraoui, A. (2020). Inner ear gene therapies take off: Current promises and future challenges. J Clin Med 9. 10.3390/jcm9072309.

Ding, B., Walton, J.P., Zhu, X., and Frisina, R.D. (2018). Age-related changes in Na, K-ATPase expression, subunit isoform selection and assembly in the stria vascularis lateral wall of mouse cochlea. Hear Res 367, 59–73. 10.1016/j.heares.2018.07.006.

Ding, J., Adiconis, X., Simmons, S.K., Kowalczyk, M.S., Hession, C.C., Marjanovic, N.D., Hughes, T.K., Wadsworth, M.H., Burks, T., Nguyen, L.T., et al. (2020). Systematic comparison of single-cell and single-nucleus RNA-sequencing methods. Nat Biotechnol 38, 737–746. 10.1038/s41587-020-0465-8.

Dobnikar, L., Taylor, A.L., Chappell, J., Oldach, P., Harman, J.L., Oerton, E., Dzierzak, E., Bennett, M.R., Spivakov, M., and Jørgensen, H.F. (2018). Disease-relevant transcriptional signatures identified in individual smooth muscle cells from healthy mouse vessels. Nat Commun 9, 4567. 10.1038/s41467-018-06891-x.

Driver, E.C., and Kelley, M.W. (2020). Development of the cochlea. Development 147. 10.1242/dev.162263.

Durán-Alonso, M.B. (2020). Stem cell-based approaches: Possible route to hearing restoration? World J Stem Cells 12, 422–437. 10.4252/wjsc.v12.i6.422.

Ebrahimi-Fakhari, D., Saffari, A., Wahlster, L., Di Nardo, A., Turner, D., Lewis, T.L., Jr., Conrad, C., Rothberg, J.M., Lipton, J.O., Kölker, S., et al. (2016). Impaired mitochondrial dynamics and mitophagy in neuronal models of tuberous sclerosis complex. Cell Rep 17, 1053–1070. 10.1016/j.celrep.2016.09.054.

Forge, A., Becker, D., Casalotti, S., Edwards, J., Marziano, N., and Nevill, G. (2003). Gap junctions in the inner ear: Comparison of distribution patterns in different vertebrates and assessement of connexin composition in mammals. J Comp Neurol 467, 207–231. 10.1002/cne.10916.

Gordon, E.J., Gale, N.W., and Harvey, N.L. (2008). Expression of the hyaluronan receptor LYVE-1 is not restricted to the lymphatic vasculature; LYVE-1 is also expressed on embryonic blood vessels. Dev Dyn 237, 1901–1909. 10.1002/dvdy.21605.

Grimaldi, A., and Tajbakhsh, S. (2021). Diversity in cranial muscles: Origins and developmental programs. Curr Opin Cell Biol 73, 110–116. 10.1016/j.ceb.2021.06.005.

Groves, A.K., and Fekete, D.M. (2012). Shaping sound in space: The regulation of inner ear patterning. Development 139, 245–257. 10.1242/dev.067074.

Hafemeister, C., and Satija, R. (2019). Normalization and variance stabilization of single-cell RNA-seq data using regularized negative binomial regression. Genome Biology 20, 296. 10.1186/s13059-019-1874-1.

Hayashi, T., Ray, C.A., and Bermingham-McDonogh, O. (2008). Fgf20 is required for sensory epithelial specification in the developing cochlea. J Neurosci 28, 5991–5999. 10.1523/jneurosci.1690-08.2008.

Homan, K.A., Gupta, N., Kroll, K.T., Kolesky, D.B., Skylar-Scott, M., Miyoshi, T., Mau, D., Valerius, M.T., Ferrante, T., Bonventre, J.V., et al. (2019). Flow-enhanced vascularization and maturation of kidney organoids in vitro. Nat Methods 16, 255–262. 10.1038/s41592-019-0325-y.

Honda, K., Kim, S.H., Kelly, M.C., Burns, J.C., Constance, L., Li, X., Zhou, F., Hoa, M., Kelley, M.W., Wangemann, P., et al. (2017). Molecular architecture underlying fluid absorption by the developing inner ear. eLife 6, e26851. 10.7554/eLife.26851.

Huebsch, N., Loskill, P., Mandegar, M.A., Marks, N.C., Sheehan, A.S., Ma, Z., Mathur, A., Nguyen, T.N., Yoo, J.C., Judge, L.M., et al. (2015). Automated video-based analysis of contractility and calcium flux in human-induced pluripotent stem cell-derived cardiomyocytes cultured over different spatial scales. Tissue Eng Part C Methods 21, 467–479. 10.1089/ten.TEC.2014.0283.

Hülse, R., Biesdorf, A., Hörmann, K., Stuck, B., Erhart, M., Hülse, M., and Wenzel, A. (2019). Peripheral vestibular disorders: An epidemiologic survey in 70 million individuals. Otol Neurotol 40, 88–95. 10.1097/MAO.0000000000002013.

Jan, T.A., Eltawil, Y., Ling, A.H., Chen, L., Ellwanger, D.C., Heller, S., and Cheng, A.G. (2021). Spatiotemporal dynamics of inner ear sensory and non-sensory cells revealed by single-cell transcriptomics. Cell Rep 36, 109358. 10.1016/j.celrep.2021.109358.

Johnson Chacko, L., Pechriggl, E.J., Fritsch, H., Rask-Andersen, H., Blumer, M.J., Schrott-Fischer, A., and Glueckert, R. (2016). Neurosensory differentiation and innervation patterning in the human fetal vestibular end organs between the gestational weeks 8-12. Front Neuroanat 10, 111. 10.3389/fnana.2016.00111.

Kaipainen, A., Korhonen, J., Mustonen, T., Van Hinsbergh, V.W., Fang, G.H., Dumont, D., Breitman, M., and Alitalo, K. (1995). Expression of the fms-like tyrosine kinase 4 gene becomes restricted to lymphatic endothelium during development. Proc Natl Acad Sci U S A 92, 3566–3570. 10.1073/pnas.92.8.3566.

Kalucka, J., De Rooij, L.P.M.H., Goveia, J., Rohlenova, K., Dumas, S.J., Meta, E., Conchinha, N.V., Taverna, F., Teuwen, L.A., Veys, K., et al. (2020). Single-cell transcriptome atlas of murine endothelial cells. Cell 180, 764–779.e720. 10.1016/j.cell.2020.01.015.

Kang, J.B., Nathan, A., Weinand, K., Zhang, F., Millard, N., Rumker, L., Moody, D.B., Korsunsky, I., and Raychaudhuri, S. (2021). Efficient and precise single-cell reference atlas mapping with Symphony. Nature Communications 12, 5890. 10.1038/s41467-021-25957-x.

Kasahara, H., Bartunkova, S., Schinke, M., Tanaka, M., and Izumo, S. (1998). Cardiac and extracardiac expression of Csx/Nkx2.5 homeodomain protein. Circ Res 82, 936–946. 10.1161/01.res.82.9.936.

Koehler, K.R., Nie, J., Longworth-Mills, E., Liu, X.P., Lee, J., Holt, J.R., and Hashino, E. (2017). Generation of inner ear organoids containing functional hair cells from human pluripotent stem cells. Nat Biotechnol 35, 583–589. 10.1038/nbt.3840.

Kolla, L., Kelly, M.C., Mann, Z.F., Anaya-Rocha, A., Ellis, K., Lemons, A., Palermo, A.T., So, K.S., Mays, J.C., Orvis, J., et al. (2020). Characterization of the development of the mouse cochlear epithelium at the single cell level. Nature Communications 11, 2389. 10.1038/s41467-020-16113-y.

Korrapati, S., Taukulis, I., Olszewski, R., Pyle, M., Gu, S., Singh, R., Griffiths, C., Martin, D., Boger, E., Morell, R.J., and Hoa, M. (2019). Single cell and single nucleus RNA-seq reveal cellular heterogeneity and homeostatic regulatory networks in adult mouse stria vascularis. Front Mol Neurosci 12, 316. 10.3389/fnmol.2019.00316.

Korsunsky, I., Millard, N., Fan, J., Slowikowski, K., Zhang, F., Wei, K., Baglaenko, Y., Brenner, M., Loh, P., and Raychaudhuri, S. (2019). Fast, sensitive and accurate integration of single-cell data with Harmony. Nature Methods 16, 1289–1296. 10.1038/s41592-019-0619-0.

Koundakjian, E.J., Appler, J.L., and Goodrich, L.V. (2007). Auditory neurons make stereotyped wiring decisions before maturation of their targets. J Neurosci 27, 14078–14088. 10.1523/jneurosci.3765-07.2007.

Lee, J., Rabbani, C.C., Gao, H., Steinhart, M.R., Woodruff, B.M., Pflum, Z.E., Kim, A., Heller, S., Liu, Y., Shipchandler, T.Z., and Koehler, K.R. (2020). Hair-bearing human skin generated entirely from pluripotent stem cells. Nature 582, 399–404. 10.1038/s41586-020-2352-3.

Lindskog, C., Linné, J., Fagerberg, L., Hallström, B.M., Sundberg, C.J., Lindholm, M., Huss, M., Kampf, C., Choi, H., Liem, D.A., et al. (2015). The human cardiac and skeletal muscle proteomes defined by transcriptomics and antibody-based profiling. BMC Genomics 16, 475. 10.1186/s12864-015-1686-y.

Locher, H., De Groot, J.C., Van Iperen, L., Huisman, M.A., Frijns, J.H., and Chuva de Sousa Lopes, S.M. (2015). Development of the stria vascularis and potassium regulation in the human fetal cochlea: Insights into hereditary sensorineural hearing loss. Dev Neurobiol 75, 1219–1240. 10.1002/dneu.22279.

Locher, H., Frijns, J.H., Van Iperen, L., De Groot, J.C., Huisman, M.A., and Chuva de Sousa Lopes, S.M. (2013). Neurosensory development and cell fate determination in the human cochlea. Neural Dev 8, 20. 10.1186/1749-8104-8-20.

Luo, X.J., Deng, M., Xie, X., Huang, L., Wang, H., Jiang, L., Liang, G., Hu, F., Tieu, R., Chen, R., and Gan, L. (2013). GATA3 controls the specification of prosensory domain and neuronal survival in the mouse cochlea. Hum Mol Genet 22, 3609–3623. 10.1093/hmg/ddt212.

MacDonald, A., Lu, B., Caron, M., Caporicci-Dinucci, N., Hatrock, D., Petrecca, K., Bourque, G., and Stratton, J.A. (2021). Single cell transcriptomics of ependymal cells across age, region and species reveals cilia-related and metal ion regulatory roles as major conserved ependymal cell functions. Front Cell Neurosci 15, 703951. 10.3389/fncel.2021.703951.

Mayordomo, R., Rodríguez-Gallardo, L., and Alvarez, I.S. (1998). Morphological and quantitative studies in the otic region of the neural tube in chick embryos suggest a neuroectodermal origin for the otic placode. J Anat 193 ( *Pt 1*), 35–48. 10.1046/j.1469-7580.1998.19310035.x.

McInturff, S., Burns, J.C., and Kelley, M.W. (2018). Characterization of spatial and temporal development of type I and type II hair cells in the mouse utricle using new cell-type-specific markers. Biol Open 7. 10.1242/bio.038083.

Mei, C., Dong, H., Nisenbaum, E., Thielhelm, T., Nourbakhsh, A., Yan, D., Smeal, M., Lundberg, Y., Hoffer, M.E., Angeli, S., et al. (2021). Genetics and the individualized therapy of vestibular disorders. Front Neurol 12, 633207. 10.3389/fneur.2021.633207.

Morsli, H., Tuorto, F., Choo, D., Postiglione, M.P., Simeone, A., and Wu, D.K. (1999). Otx1 and Otx2 activities are required for the normal development of the mouse inner ear. Development 126, 2335–2343. 10.1242/dev.126.11.2335.

Munnamalai, V., and Fekete, D.M. (2020). The acquisition of positional information across the radial axis of the cochlea. Dev Dyn 249, 281–297. 10.1002/dvdy.118.

Nishio, S.Y., Hattori, M., Moteki, H., Tsukada, K., Miyagawa, M., Naito, T., Yoshimura, H., Iwasa, Y., Mori, K., Shima, Y., et al. (2015). Gene expression profiles of the cochlea and vestibular endorgans: Localization and function of genes causing deafness. Ann Otol Rhinol Laryngol 124 *Suppl 1*, 6s–48s. 10.1177/0003489415575549.

Ohta, S., and Schoenwolf, G.C. (2018). Hearing crosstalk: The molecular conversation orchestrating inner ear dorsoventral patterning. Wiley Interdiscip Rev Dev Biol 7. 10.1002/wdev.302.

Ohyama, T., Basch, M.L., Mishina, Y., Lyons, K.M., Segil, N., and Groves, A.K. (2010). BMP signaling is necessary for patterning the sensory and nonsensory regions of the developing mammalian cochlea. J Neurosci 30, 15044–15051. 10.1523/jneurosci.3547-10.2010.

Pujol, R., and Lavigne-Rebillard, M. (1985). Early stages of innervation and sensory cell differentiation in the human fetal organ of Corti. Acta Otolaryngol Suppl 423, 43–50. 10.3109/00016488509122911.

Rau, A., Legan, P.K., and Richardson, G.P. (1999). Tectorin mRNA expression is spatially and temporally restricted during mouse inner ear development. J Comp Neurol 405, 271–280.

Roccio, M. (2021). Directed differentiation and direct reprogramming: Applying stem cell technologies to hearing research. Stem Cells 39, 375–388. 10.1002/stem.3315.

Roccio, M., and Edge, A.S.B. (2019). Inner ear organoids: New tools to understand neurosensory cell development, degeneration and regeneration. Development 146. 10.1242/dev.177188.

Roccio, M., Senn, P., and Heller, S. (2020). Novel insights into inner ear development and regeneration for targeted hearing loss therapies. Hear Res 397, 107859. 10.1016/j.heares.2019.107859.

Schupp, J.C., Adams, T.S., Cosme, C., Jr., Raredon, M.S.B., Yuan, Y., Omote, N., Poli, S., Chioccioli, M., Rose, K.A., Manning, E.P., et al. (2021). Integrated single-cell atlas of endothelial cells of the human lung. Circulation 144, 286–302. 10.1161/circulationaha.120.052318.

Shearer, A.E., Hildebrand, M.S., and Smith, R.J.H. (1993). Hereditary hearing loss and deafness overview. In GeneReviews(®), M.P. Adam, H.H. Ardinger, R.A. Pagon, S.E. Wallace, L.J.H. Bean, K.W. Gripp, G.M. Mirzaa, and A. Amemiya, eds. (University of Washington, Seattle).

Shlevkov, E., Basu, H., Bray, M.A., Sun, Z., Wei, W., Apaydin, K., Karhohs, K., Chen, P.F., Smith, J.L.M., Wiskow, O., et al. (2019). A high-content screen identifies TPP1 and aurora B as regulators of axonal mitochondrial transport. Cell Rep 28, 3224–3237.e3225. 10.1016/j.celrep.2019.08.035.

Simmons, D.D., Tong, B., Schrader, A.D., and Hornak, A.J. (2010). Oncomodulin identifies different hair cell types in the mammalian inner ear. J Comp Neurol 518, 3785–3802. 10.1002/cne.22424.

Slyper, M., Porter, C.B.M., Ashenberg, O., Waldman, J., Drokhlyansky, E., Wakiro, I., Smillie, C., Smith-Rosario, G., Wu, J., Dionne, D., et al. (2020). A single-cell and single-nucleus RNA-Seq toolbox for fresh and frozen human tumors. Nat Med 26, 792–802. 10.1038/s41591-020-0844-1.

Sontag, M.K., Yusuf, C., Grosse, S.D., Edelman, S., Miller, J.I., McKasson, S., Kellar-Guenther, Y., Gaffney, M., Hinton, C.F., Cuthbert, C., et al. (2020). Infants with congenital disorders identified through newborn screening-United States, 2015-2017. MMWR Morb Mortal Wkly Rep 69, 1265-1268. 10.15585/mmwr.mm6936a6.

Steinhart, M.R., Serdy, S.A., Van der Valk, W.H., Zhang, J., Kim, J., Lee, J., and Koehler, K.R. (2021). Defining inner ear cell type specification at single-cell resolution in a model of human cranial development. Cell Reports, Sneak Peek.

Stoeckius, M., Zheng, S., Houck-Loomis, B., Hao, S., Yeung, B.Z., Mauck, W.M., Smibert, P., and Satija, R. (2018). Cell hashing with barcoded antibodies enables multiplexing and doublet detection for single cell genomics. Genome Biology 19, 224. 10.1186/s13059-018-1603-1.

Stojkovic, M., Han, D., Jeong, M., Stojkovic, P., and Stankovic, K.M. (2021). Human induced pluripotent stem cells and CRISPR/Cas-mediated targeted genome editing: Platforms to tackle sensorineural hearing loss. Stem Cells 39, 673–696. 10.1002/stem.3353.

Taiber, S., and Avraham, K.B. (2019). Genetic therapies for hearing loss: Accomplishments and remaining challenges. Neurosci Lett 713, 134527. 10.1016/j.neulet.2019.134527.

Tang, P.C., Hashino, E., and Nelson, R.F. (2020). Progress in modeling and targeting inner ear disorders with pluripotent stem cells. Stem Cell Reports 14, 996–1008. 10.1016/j.stemcr.2020.04.008.

Truong, K., Ahmad, I., Jason Clark, J., Seline, A., Bertroche, T., Mostaert, B., Van Daele, D.J., and Hansen, M.R. (2018). Nf2 mutation in Schwann cells delays functional neural recovery following injury. Neuroscience 374, 205–213. 10.1016/j.neuroscience.2018.01.054.

Van Beeck Calkoen, E.A., Engel, M.S.D., Van de Kamp, J.M., Yntema, H.G., Goverts, S.T., Mulder, M.F., Merkus, P., and Hensen, E.F. (2019). The etiological evaluation of sensorineural hearing loss in children. Eur J Pediatr 178, 1195–1205. 10.1007/s00431-019-03379-8.

Van Beelen, E.S.A., Van der Valk, W.H., De Groot, J., Hensen, E.F., Locher, H., and Van Benthem, P.P.G. (2020). Migration and fate of vestibular melanocytes during the development of the human inner ear. Dev Neurobiol 80, 411–432. 10.1002/dneu.22786.

Van der Valk, W.H., Steinhart, M.R., Zhang, J., and Koehler, K.R. (2021). Building inner ears: Recent advances and future challenges for in vitro organoid systems. Cell Death Differ 28, 24–34. 10.1038/s41418-020-00678-8.

Vandekerckhove, J., Bugaisky, G., and Buckingham, M. (1986). Simultaneous expression of skeletal muscle and heart actin proteins in various striated muscle tissues and cells. A quantitative determination of the two actin isoforms. J Biol Chem 261, 1838–1843.

Wang, A., Shearer, A.E., Zhou, G.W., Kenna, M., Poe, D., Licameli, G.R., and Brodsky, J.R. (2021). Peripheral vestibular dysfunction is a common occurrence in children with non-syndromic and syndromic genetic hearing loss. Front Neurol 12, 714543. 10.3389/fneur.2021.714543.

Wangzhou, A., Paige, C., Ray, P.R., Dussor, G., and Price, T.J. (2021). Diversity of receptor expression in central and peripheral mouse neurons estimated from single cell RNA sequencing. Neuroscience 463, 86–96. 10.1016/j.neuroscience.2021.03.017.

Whalen, R.G., Butler-Browne, G.S., and Gros, F. (1978). Identification of a novel form of myosin light chain present in embryonic muscle tissue and cultured muscle cells. J Mol Biol 126, 415–431. 10.1016/0022-2836(78)90049-9.

White, R.M., and Zon, L.I. (2008). Melanocytes in development, regeneration, and cancer. Cell Stem Cell 3, 242–252. 10.1016/j.stem.2008.08.005.

WHO (2021). Deafness and Hearing Loss. https://www.who.int/news-room/fact-sheets/detail/deafness-and-hearing-loss.

Wilkerson, B.A., Zebroski, H.L., Finkbeiner, C.R., Chitsazan, A.D., Beach, K.E., Sen, N., Zhang, R.C., and Bermingham-McDonogh, O. (2021). Novel cell types and developmental lineages revealed by single-cell RNA-seq analysis of the mouse crista ampullaris. Elife 10. 10.7554/eLife.60108.

Wilson, P.A., and Hemmati-Brivanlou, A. (1995). Induction of epidermis and inhibition of neural fate by Bmp-4. Nature 376, 331–333. 10.1038/376331a0.

Wu, C.L., Dicks, A., Steward, N., Tang, R., Katz, D.B., Choi, Y.R., and Guilak, F. (2021). Single cell transcriptomic analysis of human pluripotent stem cell chondrogenesis. Nat Commun 12, 362. 10.1038/s41467-020-20598-y.

Yamamoto, R., Ohnishi, H., Omori, K., and Yamamoto, N. (2021). In silico analysis of inner ear development using public whole embryonic body single-cell RNA-sequencing data. Dev Biol 469, 160–171. 10.1016/j.ydbio.2020.10.009.

Yu, K.S., Frumm, S.M., Park, J.S., Lee, K., Wong, D.M., Byrnes, L., Knox, S.M., Sneddon, J.B., and Tward, A.D. (2019). Development of the mouse and human cochlea at single cell resolution. bioRxiv, 739680. 10.1101/739680.

Zeisel, A., Hochgerner, H., Lönnerberg, P., Johnsson, A., Memic, F., Van der Zwan, J., Häring, M., Braun, E., Borm, L.E., La Manno, G., et al. (2018). Molecular architecture of the mouse nervous system. Cell 174, 999–1014.e1022. 10.1016/j.cell.2018.06.021.

Zine, A., Messat, Y., and Fritzsch, B. (2021). A human induced pluripotent stem cell-based modular platform to challenge sensorineural hearing loss. Stem Cells 39, 697–706. 10.1002/stem.3346.

