## Supplementary material for "A Single-Cell Level Comparison of Human Inner Ear Organoids and the Human Cochlea and Vestibular Organs": Table S1. Evaluation of 158 SNHL genes and their expression patterns in IEOs.

| GENE | OMIM | NONSYNDROMIC HL | SYNDROMIC HL | EXPRESSION DESCRIBED IN LITERATURE | VESTIBULAR PHENOTYPE REPORTED | SCORED IEO RNA EXPRESSION | IHC CONFIRMED | NOTE | REFERENCES |
|---|---|---|---|---|---|---|---|---|---|
| ACTG1 | 102560 | DFNA20/26 | | Neurosensory epithelia | | 1 | | | Zhu et al., 2003; van Wijk et al., 2003; Perrin et al., 2010 |
| ADCY1 | 103072 | DFNB44 | | Neurosensory epithelia | | 3 | | | Santos-Cortez et al., 2014 |
| ADGRV1 | 602851 | | Usher Syndrome | Hair cell (stereocilia) | Yes | 5 | | | Weston et al., 2004 |
| AIFM1 | 300169 | DFNX5 | | Multiple cell types | Yes | 4 | | | Zong et al., 2015; Wang et al., 2020 |
| ATP2B2 | 108733 | DFNA73 | | Hair cell (stereocilia) | | 5 | | | Smits et al., 2019; Hill et al., 2006 |
| BDP1 | 607012 | DFNB49 | | Multiple cell types | | 3 | | | Girotto et al., 2013 |
| BSND | 606412 | DFNB73 | Bartter Syndrome | Stria vascularis, dark cell epithelium | Yes | 1 | Yes | | Riazuddin et al., 2009; Rickheit et al., 2008 |
| CABP2 | 607314 | DFNB93 | | Hair cell, spiral ganglion | | 5 | | | Schrauwen et al., 2012; Oestreicher et al., 2021 |
| CCDC50 | 611051 | DFNA44 | | Multiple cell types | | 2 | | Involved in early inner ear formation | Modamio-Hoybjor et al., 2007 |
| CD164 | 603356 | DFNA66 | | Multiple cell types | | 4 | | | Nyegaard et al., 2015 |
| CDC14A | 603504 | DFNB32/105 | | Hair cell (stereocilia) | | 2 | | | Delmaghani et al., 2016; Imtiaz et al., 2017 |
| CDH23 | 605516 | DFNB12 | Usher Syndrome | Hair cell, Reissner's membrane | Yes | 5 | Yes | | Bork et al., 2001; Bolz et al., 2001; Wafa et al., 2020; Siemens et al., 2004 |
| CEACAM16 | 614591 | DFNA4B; DFNB108 | | Neurosensory epithelia | | 0 | | Expression not shown for specific cell types | Zheng et al., 2011; Booth et al., 2018 |
| CHD7 | 608892 | | CHARGE Syndrome | Multiple cell types | Yes | 4 | | | Vissers et al., 2004; Abadie et al., 2000; Ahmed et al., 2021 |
| CIB2 | 605564 | DFNB48 | | Neurosensory epithelia | Yes | 4 | | | Riazuddin et al., 2012; Michel et al., 2017 |
| CLDN14 | 605608 | DFNB29 | | Multiple cell types | | 4 | | | Wilcox et al., 2001 |
| CLDN9 | 615799 | DFNB108 | | Neurosensory epithelia | | 4 | | | Sineni et al., 2019 |
| CLIC5 | 607293 | DFNB103 | | Neurosensory epithelia | Yes | 5 | | | Seco et al., 2014; Gagnon et al., 2006 |
| CLPP | 601119 | | Perrault Syndrome | Neurosensory epithelia | | 3 | | | Jenkinson et al., 2013; Forli et al., 2021 |
| CLRN1 | 606397 | | Usher Syndrome | Hair cell, spiral ganglion | Yes | 5 | | | Joensuu et al., 2001; Zallochi et al., 2010 |
| CLRN2 | 618988 | DFNB108 | | Hair cell (stereocilia) | | 4 | | | Vona et al., 2021; Dunbar et al., 2019 |
| COCH | 603196 | DFNA9 | | POM | Yes | 5 | | | Robertson et al., 1998; Kim et al., 2016 |

| | | | | | | | | | |
|---|---|---|---|---|---|---|---|---|---|
| COL11A1 | 120280 | DFNA37 | Stickler Syndrome | Multiple cell types | Yes | 4 | | | Booth et al., 2018; Richards et al., 1996; Acke et al., 2012; Shpargel et al., 2009 |
| COL11A2 | 120290 | DFNA13; DFNB53 | Stickler Syndrome | Multiple cell types | Yes | 4 | | | McGuirt et al., 1999; Chen et al., 2005; Vikkula et al., 1995; Acke et al., 2012; Shpargel et al., 2009 |
| COL2A1 | 120140 | | Stickler Syndrome | Multiple cell types | Yes | 4 | | | Ahmad et al., 1991; Acke et al., 2012; Khetarpal et al., 1994 |
| COL4A3 | 120070 | | Alport Syndrome | Multiple cell types | Yes | 3 | | | Mochizuki et al., 1994; Barozzi et al., 2020; Zhender et al., 2005 |
| COL4A4 | 120131 | | Alport Syndrome | Multiple cell types | Yes | 3 | | | Mochizuki et al., 1994; Barozzi et al., 2020; Zhender et al., 2005 |
| COL4A5 | 303630 | | Alport Syndrome | Multiple cell types | Yes | 4 | | | Barker et al., 1990; Barozzi et al., 2020; Zhender et al., 2005 |
| COL4A6 | 303631 | DFNX6 | | Multiple cell types | | 4 | | | Rost et al., 2014 |
| COL9A1 | 120210 | | Stickler Syndrome | Multiple cell types | Yes | 4 | | | Van Camp et al., 2006; Acke et al., 2012; Sivakumaran et al., 2006 |
| COL9A2 | 120260 | | Stickler Syndrome | Multiple cell types | Yes | 4 | | | Baker et al., 2011; Acke et al., 2012; Chacko et al., 2021 |
| CRYM | 123740 | DFNA40 | | POM | | 5 | | | Abe et al., 2003 |
| DCDC2 | 605755 | DFNB66 | | Neurosensory epithelia | | 3 | | | Grati et al., 2015 |
| DIABLO | 605219 | DFNA64 | | Hair cell | | 1 | | | Cheng et al., 2011 |
| DIAPH1 | 602121 | DFNA1 | | Neurosensory epithelia | | 3 | | | Lynch et al., 1997; Ninoyu, 2020 |
| DIAPH3 | 614567 | AUNA1 | | Unknown | | 3 | | | Kim et al., 2004; Schoen et al., 2010 |
| DMXL2 | 612186 | DFNA73 | | Hair cell | | 3 | | | Chen et al., 2016 |
| EDN3 | 131242 | | Waardenburg Syndrome | Multiple cell types | Yes | 3 | | Function known to be related to melanocytes | Edery et al., 1996 |
| EDNRB | 131244 | | Waardenburg Syndrome | Melanocytes | Yes | 4 | | Function known to be related to melanocytes | Attié et al., 1995 |
| ELMOD3 | 615427 | DFNB88 | | Neurosensory epithelia | | 3 | | | Jaworek et al., 2013; Li et al., 2019 |
| EPS8 | 600206 | DFNB102 | | Hair cell (stereocilia) | | 2 | | | Behlouli et al., 2014 |
| EPS8L2 | 614988 | DFNB106 | | Hair cell (stereocilia) | | 3 | | | Dahmani et al., 2015; Furness et al., 2013 |
| ERAL1 | 607435 | | Perrault Syndrome | Multiple cell types | | 2 | | | Chatzispyrou et al., 2017 |
| ESPN | 606351 | DFNB36 | | Hair cell (stereocilia) | Yes | 4 | | | Naz et al., 2004 |
| ESRP1 | 612959 | DFNB108 | | Multiple cell types | Yes | 2 | | | Rohacek et al., 2017 |
| ESRRB | 602167 | DFNB35 | | Multiple cell types | | 4 | | | Collin et al., 2008 |
| EYA1 | 601653 | | BOR Syndrome | Multiple cell types | Yes | 3 | | Involved in early inner ear formation | Abdelhak et al., 1997 |

| | | | | | | | | | |
|---|---|---|---|---|---|---|---|---|---|
| EYA4 | 603550 | DFNA10 | | Multiple cell types | | 3 | | Involved in early inner ear formation | Wayne et al., 2001; Matsuzaki et al., 2018 |
| FOXI1 | 601093 | | Pendred Syndrome | Endolymphatic epithelia | Yes | 0 | | | Yang et al., 2007; Vidarsson et al., 2009 |
| GAB1 | 604439 | DFNB26 | | Multiple cell types | | 3 | | | Yousaf et al., 2018 |
| GAS2 | 602835 | DFNB108 | | Supporting cells | | 5 | | | Chen et al., 2021 |
| GIPC3 | 608792 | DFNB15/72/95 | | Neurosensory epithelia | | 5 | | | Ain et al., 2007; Rehman et al., 2011; Charizopoulou et al., 2011 |
| GJB2 | 121011 | DFNA3A, DFNB1A | | Multiple cell types | Yes | 4 | Yes | | Kelsell et al., 1997; Dodson et al., 2011 |
| GJB3 | 603324 | DFNA2B | | POM | | 1 | | | Xia et al., 1998; Lopez-Bigas et al., 2001 |
| GJB6 | 604418 | DFNA3B | | Multiple cell types | Yes | 4 | Yes | | Grifa et al., 1999 |
| GPSM2 | 609245 | DFNB82 | | Multiple cell types | | 4 | | | Walsh et al., 2010 |
| GRAP | 604330 | DFNB108 | | Vestibulocochlear ganglion | | N/A | | | Li et al., 2019 |
| GRHL2 | 608576 | DFNA28 | | Multiple cell types | | 3 | | | Peters et al., 2002; Hosoya et al., 2016 |
| GRXCR1 | 613283 | DFNB25 | | Hair cell (stereocilia) | Yes | 5 | | | Schraders et al., 2010 |
| GRXCR2 | 615762 | DFNB101 | | Hair cell (stereocilia) | Yes | 4 | | | Imtiaz et al., 2014; Avenarius et al., 2018 |
| GSDME | 608798 | DFNA5 | | Unknown | Yes | 3 | | | Van Laer et al., 1998 |
| HARS1 | 142810 | | Usher Syndrome | Hair cell | Yes | 0 | | | Puffenberger et al., 2012; Castiglione et al., 2022 |
| HARS2 | 600783 | | Perrault Syndrome | Multiple cell types | | 2 | | | Pierce et al., 2011 |
| HGF | 142409 | DFNB39 | | Stria vascularis | | 1 | | | Schultz et al., 2009 |
| HOMER2 | 604799 | DFNA68 | | Hair cell (stereocilia) | | 4 | | | Azaiez et al., 2015 |
| HSD17B4 | 601860 | | Perrault Syndrome | Multiple cell types | | 3 | | | Pierce et al., 2011 |
| IFNLR1 | 607404 | DFNA2C | | Multiple cell types | | 3 | | | Gao et al., 2017 |
| ILDR1 | 609739 | DFNB42 | | Multiple cell types | | 3 | | | Borck et al., 2011 |
| KARS1 | 601421 | DFNB89 | | Neurosensory epithelia | | 0 | | | Santos-Cortez et al., 2013 |
| KCNE1 | 176261 | | Jervell & Lange-Nielsen Syndrome | Stria vascularis, dark cell epithelium | Yes | 1 | | Expressed specifically in nonsensory epithelial | Tyson et al., 1997; Schulze-Bahr et al., 1997 |
| KCNJ10 | 602208 | | Pendred Syndrome | Stria vascularis, dark cell epithelium | Yes | 2 | | | Yang et al., 2009; Jin et al., 2006; Locher et al., 2015 |
| KCNQ4 | 603537 | DFNA2A | | Hair cell | | 2 | | | Kubisch et al., 1999; Kharkovets et al., 2020; Smith et al., 2018 |
| KITLG | 184745 | DFNA69 | Waardenburg Syndrome | Multiple cell types | Yes | 1 | | Function known to be related to melanocytes | Zazo Seco et al., 2015; van Beelen et al., 2020 |

| | | | | | | | | | |
|---|---|---|---|---|---|---|---|---|---|
| KNCQ1 | 607542 | | Jervell & Lange-Nielsen Syndrome | Stria vascularis, dark cell epithelium | Yes | 0 | | Expressed specifically in nonsensory epithelial | Neyroud et al., 1997 |
| LARS2 | 604544 | | Perrault Syndrome | Multiple cell types | | 3 | | | Pierce et al., 2013; Xu et al., 2021 |
| LHFPL5 | 609427 | DFNB66/67 | | Neurosensory epithelia | | 5 | | | Tlili et al., 2005; Shabbir et al., 2006; Kalay et al., 2006 |
| LMX1A | 600298 | DFNA7 | | Multiple cell types | Yes | 4 | | Involved in early inner ear formation | Wesdorp et al., 2018; Chizhikov et al., 2021 |
| LOXHD1 | 613072 | DFNB77 | | Hair cell | | 5 | | | Grillet et al., 2009 |
| LRTOMT | 612414 | DFNB63 | | Neurosensory epithelia | | 2 | | | Ahmed et al., 2008; Du et al., 2008 |
| MAP1B | 157129 | DFNA78 | | Multiple cell types | | 3 | | | Cui et al., 2020 |
| MARVELD2 | 610572 | DFNB49 | | Multiple cell types | | 4 | | | Riazuddin et al., 2006 |
| MCM2 | 116945 | DFNA70 | | Multiple cell types | | 1 | | | Gao et al., 2015 |
| MET | 164860 | DFNB97 | | Multiple cell types | | 1 | | | Mujtaba et al., 2015 |
| METTL13 | 617987 | DFNM1 | | Multiple cell types | | 0 | | | Riazuddin et al., 2000; Yousaf et al., 2018 |
| MIR96 | 611606 | DFNA50 | | Hair cell, spiral ganglion | | 0 | | | Mencia et al., 2009; Ushakov et al., 2013; Lewis et al., 2009 |
| MITF | 156845 | | Waardenburg Syndrome | Melanocytes | Yes | 5 | | Function known to be related to melanocytes | Tassabehji et al., 1994 |
| MPZL2 | 604873 | DFNB108 | | Multiple cell types | | 3 | | | Wesdorp et al., 2018 |
| MSRB3 | 613719 | DFNB74 | | Neurosensory epithelia | | 4 | | | Waryah et al., 2009; Ahmed et al., 2011 |
| MYH14 | 608568 | DFNA4A | | Multiple cell types | | 2 | | | Donaudy et al., 2004 |
| MYH9 | 160775 | DFNA17 | | Multiple cell types | | 2 | | | Lalwani et al., 2000; Mhatre et al., 2006 |
| MYO15A | 602666 | DFNB3 | | Hair cell (stereocilia) | Yes | 5 | | | Wang et al., 1998 |
| MYO3A | 606808 | DFNA73; DFNB30 | | Hair cell (stereocilia) | | 5 | | | Grati et al., 2016; Walsh et al., 2002 |
| MYO6 | 600970 | DFNA22; DFNB37 | | Hair cell | Yes | 5 | | | Melchionda et al., 2001; Ahmed et al., 2003 |
| MYO7A | 276903 | DFNA11; DFNB2 | Usher Syndrome | Hair cell | Yes | 5 | Yes | | Liu et al., 1997; Weil et al., 1997; Wafa et al., 2020 |
| NARS2 | 612803 | DFNB94 | | Multiple cell types | | 0 | | | Simon et al., 2015 |
| NDP | 300658 | | Norrie Disease | Stria vascularis | | 0 | | | Berger et al., 1992; Chen et al., 1992; Hayashi et al., 2021 |
| NLRP3 | 606416 | DFNA34 | | Macrophages | | N/A | | | Nakanishi et al., 2017 |
| OSBPL2 | 606731 | DFNA67 | | Hair cell (stereocilia) | | 2 | | | Xing et al., 2014; Thoenes et al., 2015 |
| OTOA | 607038 | DFNB22 | | Multiple cell types | | 3 | | | Zwaenepoel et al., 2002; Lukashkin et al., 2012 |

| | | | | | | | | | |
|---|---|---|---|---|---|---|---|---|---|
| OTOF | 603681 | DFNB9 | | Hair cell, spiral ganglion | | 5 | Yes | | Yasunaga et al., 1999; Santarelli et al., 2021 |
| OTOG | 604487 | DFNB18B | | Multiple cell types | Yes | 3 | | | Schraders et al., 2012 |
| OTOGL | 614925 | DFNB84 | | Multiple cell types | Yes | 4 | | | Yariz et al., 2012 |
| P2RX2 | 600844 | DFNA41 | | Multiple cell types | Yes | 1 | | | Yan et al., 2013; Jarlebark et al., 2002 |
| PAX3 | 606597 | | Waardenburg Syndrome | Melanocytes | Yes | 5 | | Function known to be related to melanocytes | Tassebehji et al., 1992; Zlotogora et al., 1995; Kim et al., 2014 |
| PCDH15 | 605514 | DFNB23 | Usher Syndrome | Hair cell (stereocilia) | Yes | 5 | | | Ahmed et al., 2003; Ahmed et al., 2001, Alagramam et al., 2001; Webb et al., 2011; Wafa et al., 2020 |
| PDE1C | 602987 | DFNA73 | | Hair cell | | 2 | | | Wang et al., 2018 |
| PDZD7 | 612971 | DFNB57 | | Hair cell | Yes | 5 | | | Booth et al, 2015; Chen et al., 2014 |
| PJVK | 610219 | DFNB59 | | Hair cell | | 4 | | | Delmaghani et al., 2006; Kazmierczak et al., 2017 |
| PLS1 | 602734 | DFNA73 | | Hair cell (stereocilia) | | 4 | | | Morgan et al., 2019; Krey et al., 2016 |
| PNPT1 | 610316 | DFNB70 | | Multiple cell types | | 3 | | | von Ameln et al., 2012 |
| POLR1C | 610060 | | Treacher Collins Syndrome | Craniofacial neural crest | Yes | 2 | | Involved in early inner ear formation | Dauwerse et al., 2010 |
| POLR1D | 613715 | | Treacher Collins Syndrome | Craniofacial neural crest | Yes | 2 | | Involved in early inner ear formation | Dauwerse et al., 2010 |
| POU3F4 | 300039 | DFNX2 | | POM | Yes | 5 | Yes | | De Kok et al., 1995; Song et al., 2012; Vore et al., 2005 |
| POU4F3 | 602460 | DFNA15 | | Hair cell | Yes | 5 | | | Vahava et al., 1998, van Drunen et al., 2009 |
| PPIP5K2 | 611648 | DFNB100 | | Multiple cell types | | 3 | | | Yousaf et al., 2018 |
| PRPS1 | 311850 | DFNX1 | | Hair cell, spiral ganglion | | 2 | | | Liu et al., 2010 |
| PTPRQ | 603317 | DFNA73; DFNB84 | | Hair cell | Yes | 5 | | | Eisenberger et al., 2017; Schraders et al., 2010; Goodyear et al., 2003 |
| RDX | 179410 | DFNB24 | | Hair cell | Yes | 2 | | | Khan et al., 2007; Kitajiri et al., 2004 |
| REST | 600571 | DFNA27 | | Unknown | | 2 | | | Nakano et al., 2018 |
| RIPOR2 | 611410 | DFNA78; DFNB104 | | Hair cell | | 5 | | | de Bruijn et al., 2020; Diaz-Horta et al., 2019; Diaz-Horta et al., 2014 |
| ROR1 | 602336 | DFNB108 | | Multiple cell types | | 2 | | | Diaz-Horta et al., 2016 |
| S1PR2 | 605111 | DFNB68 | | Multiple cell types | Yes | 2 | | | Santos-Cortez et al., 2016; Ingham et al., 2016 |
| SCD5 | 608370 | DFNA73 | | Unknown | | 3 | | | Lu et al., 2020 |
| SEMA3E | 608166 | | CHARGE Syndrome | Unknown | Yes | 3 | | | Lalani et al., 2004; Abadie et al., 2000 |

| | | | | | | | | | |
|---|---|---|---|---|---|---|---|---|---|
| SERPINB6 | 173321 | DFNB91 | | Hair cell | | 2 | | | Sirmaci et al., 2010 |
| SIX1 | 601205 | DFNA23 | BOR Syndrome | Hair cell | Yes | 3 | Yes | Involved in early inner ear formation | Mosrati et al., 2011; Ruf et al., 2003 |
| SIX5 | 600963 | | BOR Syndrome | Early development | Yes | 2 | | Involved in early inner ear formation | Hoskins et al., 2007 |
| SLC12A2 | 600840 | DFNA78 | | Stria vascularis | Yes | 2 | | | Mutai et al., 2020 |
| SLC17A8 | 607557 | DFNA25 | | Hair cell, spiral ganglion | | 4 | | | Ruel et al., 2008 |
| SLC22A4 | 604190 | DFNB60 | | Endothelial cells | | N/A | | | Ben Said et al., 2016 |
| SLC26A4 | 605646 | DFNB4 | Pendred Syndrome | Multiple cell types | Yes | 3 | Yes | | Li et al., 1998; Everett et al., 1997 |
| SLC26A5 | 604943 | DFNB61 | | Hair cell | | 4 | | | Liu et al., 2003 |
| SMPX | 300226 | DFNX4 | | Multiple cell types | | 4 | | | Schraders et al., 2011; Huebner et al., 2011; Tu et al., 2021 |
| SNAI2 | 602150 | | Waardenburg Syndrome | Unknown | Yes | 3 | | | Sánchez-Martin et al., 2002 |
| SOX10 | 602229 | | Waardenburg Syndrome | Multiple cell types | Yes | 5 | Yes | Function known to be related to melanocytes | Bondurand et al., 2007; Pingault et al., 1998; Black et al., 2011 |
| SPNS2 | 612584 | DFNB108 | | Multiple cell types | Yes | 3 | | | Chen et al., 2014; Ingham et al., 2019 |
| STRC | 606440 | DFNB16 | | Hair cell (stereocilia) | Yes | 5 | | | Verpy et al., 2001; Dodson et al., 2011 |
| SYNE4 | 615535 | DFNB76 | | Hair cell, spiral ganglion | | 3 | | | Horn et al., 2013; Taiber et al., 2021 |
| TBC1D24 | 613577 | DFNA65; DFNB86 | | Hair cell, spiral ganglion | | 2 | | | Azaiez et al., 2014; Zhang et al., 2014; Rehman et al., 2014; Ozieblo et al. 2021 |
| TCOF1 | 606847 | | Treacher Collins Syndrome | Craniofacial neural crest | Yes | 0 | | Involved in early inner ear formation | Dixon et al., 1996 |
| TECTA | 602574 | DFNA8/12; DFNB21 | | Tectorial membrane/otolith membrane | | 5 | | | Verhoeven et al., 1998; Mustapha et al., 1999; Street et al., 2008 |
| TJP2 | 607709 | DFNA51 | | Neurosensory epithelia | | 4 | | | Walsh et al., 2010 |
| TMC1 | 606706 | DFNA36; DFNB7/11 | | Multiple cell types | Yes | 4 | | | Kurima et al., 2002 |
| TMEM132E | 616178 | DFNB99 | | Multiple cell types | | 0 | | | Li et al., 2015; Liaqat et al., 2019 |
| TMIE | 607237 | DFNB6 | | Multiple cell types | Yes | 3 | | | Naz et al., 2002; Zhao et al., 2014 |
| TMPRSS3 | 605511 | DFNB8/10 | | Multiple cell types | Yes | 4 | | | Scott et al., 2001; Fasquelle et al., 2011 |
| TNC | 187380 | DFNA56 | | Hair cell | | 5 | | | Zhao et al., 2013; Son et al., 2012 |

| | | | | | | | | | |
|---|---|---|---|---|---|---|---|---|---|
| TPRN | 613354 | DFNB79 | | Multiple cell types | | 2 | | | Rehman et al., 2010; Li et al., 2010 |
| TRIOBP | 609761 | DFNB28 | | Neurosensory epithelia | | 3 | | | Shahin et al., 2006; Riazuddin et al., 2006; Babahosseini et al., 2021 |
| TRRAP | 603015 | DFNA73 | | Unknown | | 2 | | | Xia et al., 2019 |
| TSPEAR | 612920 | DFNB98 | | Multiple cell types | | 1 | | | Delmaghani et al., 2012 |
| TWNK | 606075 | | Perrault Syndrome | Vestibulocochlear ganglion | | N/A | | | Morino et al., 2014 |
| USH1C | 605242 | DFNB18 | Usher Syndrome | Hair cell (stereocilia) | Yes | 4 | | | Ouyang et al., 2002; Ahmed et al., 2002; Verpy et al., 2000; Wafa et al., 2020 |
| USH1G | 607696 | | Usher Syndrome | Hair cell (stereocilia) | Yes | 1 | | | Weil et al., 2003; Wafa et al., 2020 |
| USH2A | 608400 | | Usher Syndrome | Hair cell (stereocilia) | Yes | 5 | | | Eudy et al., 1998; Wafa et al, 2021 |
| WBP2 | 606962 | DFNB108 | | Multiple cell types | | 3 | | | Buniello et al., 2016 |
| WFS1 | 606201 | DFNA6/14/38 | | Multiple cell types | | 3 | | | Bespalova et al., 2001; Young et al., 2001; Cryns et al., 2003; Bramhall et al., 2008 |
| WHRN | 607928 | DFNB31 | Usher Syndrome | Hair cell (stereocilia) | Yes | 4 | Yes | | Mburu et al., 2003; Ebermann et al., 2007; Mathur et al., 2019; Stemerdink et al., 2022 |

**Table S1. Evaluation of 158 SNHL genes and their expression patterns in IEOs.** Genes are related to nonsyndromic and/or a selection of syndromic causes of SNHL. Expression within inner ear cell types and presence of vestibular complaints/abnormalities are described using available literature. RNA expression within the IEO is rated 0-5 or N/A (non-applicable) if the expression is outside the hair cells, otic epithelium and POM. Rating 0: No expression observed; Rating 1: Sparse expression observed; Rating 2: Some expression observed and nonspecific; Rating 3: Expression observed either nonspecific or not in accordance with literature; Rating 4: Sparse expression observed in accordance with literature; Rating 5: Expression specific and accordance with literature. Genes that are rated “4” or “5” are highlighted.
